# High temperature-induced diapause transiently primes progeny for dauer formation in *C. elegans*

**DOI:** 10.64898/2026.08.26.747381

**Authors:** Esteban Retamales, James Lee, Andrea Calixto

## Abstract

Environmental stress during early development can have lasting effects on reproduction and developmental plasticity in *Caenorhabditis elegans*. Here, we compared the consequences of two dauer-inducing stressors—high temperature and crowding—on fertility, dauer formation, and intergenerational gene expression. Entry into the dauer stage protected animals from stress-induced sterility, with high-temperature–induced diapause (HID) providing strong preservation of reproductive capacity. Remarkably, the progeny of temperature-induced post-dauers (*PD-temp*) displayed a twofold increase in dauer formation upon re-exposure to heat, revealing a transient intergenerational enhancement of HID. This effect was stimulus-specific, as parental heat exposure suppressed pheromone-induced dauer formation in progeny, while parental pheromone exposure did not enhance HID. This increased dauer propensity was reset after a single stress-free generation. RNA-seq across three generations identified a transient F1-specific gene expression signature associated with enhanced dauer formation upon re-exposure to heat. Functional analyses showed that *snpc-1.3*, F49F1.7, and Y69A2AR.12 promote HID. In parallel, *vit-3* expression was selectively reduced in F1 progeny of *PD-temp* animals, and *vit-3* mutants exhibited increased dauer formation at 27°C, suggesting that *vit-3* normally restrains HID. Consistent with previous work from our group implicating RNAi pathways in environmentally induced diapause and inherited stress responses, we find that endogenous RNAi pathways also modulate HID across generations. Multiple RNAi pathway components contributed to HID, while the nuclear RNAi factor *nrde-2* was specifically required for the intergenerational increase in dauer formation. Tissue-specific rescue experiments further suggest that coordinated RNAi activity across tissues contributes differently to parental HID and progeny responses. Together, these findings identify HID as a distinct stress-induced developmental program that transiently modifies progeny responses to recurring thermal stress while preserving reproductive fitness. Our results further indicate that the physiological and intergenerational consequences of dauer entry depend on the environmental cue that induces diapause.

**Condensed abstract:** Environmental stressors such as heat and crowding promote entry into diapause in the nematode *C. elegans*. We show that the progeny of temperature-induced post-dauers exhibit a twofold increase in dauer formation upon re-exposure to heat, revealing a transient intergenerational enhancement of high-temperature–induced diapause (HID) that resets after a single stress-free generation.

To investigate the molecular basis of this transient intergenerational enhancement of HID, we combined transcriptomic and functional analyses and identified genes that promote HID, including *snpc-1.3*, F49F1.7, and Y69A2AR.1, while *vit-3* acts as a negative regulator of HID and is selectively downregulated in progeny derived from temperature-induced post-dauers. Consistent with previous work implicating RNAi pathways in inherited stress responses, we further show that the nuclear RNAi factor NRDE-2 is required for the intergenerational increase in dauer formation. In parallel, entry into diapause protected animals from stress-induced sterility, with HID preserving reproductive capacity across generations. Together, these findings identify HID as a reversible developmental program that transiently modifies progeny responses to recurring thermal stress while buffering the detrimental effects of heat stress on fertility.

## Introduction

In *Caenorhabditis elegans*, stress-induced plasticity influences developmental decisions: Exposure to pathogens, for example, induces entry into diapause (*Palominos et al.*, 2017). These responses require effectors of the RNA interference (RNAi) pathway at multiple levels, as well as specific small RNAs from the host (*Gabaldón et al.*, 2020) or bacteria (*Legüe et al.*, 2022), together with chromatin-based mechanisms such as histone modifications (*Legüe et al.*, 2022). Extended starvation during dauer diapause induces changes in starvation resistance, lifespan, and gene expression plasticity across generations, demonstrating that dauer-associated nutritional stress can have persistent physiological consequences (*Webster et al.*, 2018).

Dauer diapause is metabolically analogous to mammalian hibernation and is characterized by arrested growth and reproduction (*Cassada and Russell*, 1975). Dauer larvae can survive for months without food and tolerate extreme desiccation and freezing (*Erkut et al.*, 2016; *Shatilovich et al.*, 2023). In addition to promoting survival, dauer formation facilitates dispersal through nictation, a specialized behavior that allows dauers to hitchhike on carrier animals to colonize new habitats (*Lee et al.*, 2011). Multiple environmental cues induce dauer formation, including food scarcity (*Cassada and Russell*, 1975), elevated pheromone levels associated with crowding (*Golden and Riddle*, 1982), exposure to pathogens (*Palominos et al.*, 2017), high temperature (≥27 °C) (*Ailion and Thomas*, 2000), and interactions with natural microbial consortia (*Serey et al.*, 2025). Notably, *daf-22* mutants deficient in pheromone biosynthesis can still form dauers at 27 °C in the absence of exogenous pheromone (*Ailion and Thomas*, 2000), indicating that temperature alone is sufficient to trigger this developmental switch.

Passage through pheromone-induced dauer leaves lasting signatures on adult physiology, including changes in gene expression, fertility, and lifespan (*Hall et al.*, 2010). However, it remains unknown whether dauer entry itself functions as a developmental experience that alters future diapause decisions.

Specifically, it is unclear whether prior dauer passage in the parental generation influences the magnitude of dauer formation in subsequent generations—that is, the proportion of individuals within a population that commit to diapause upon re-exposure to inducing cues.

Here, we test whether parental experience of dauer diapause biases dauer propensity and reproductive fitness across generations. Using high-temperature-induced diapause (HID) and crowding-induced dauer as parallel paradigms, we ask whether prior dauer passage alters developmental decision-making upon re-exposure to heat stress. We combine phenotypic assays with RNA sequencing across three generations—naïve P0, re-exposed F1, and rested F2— to dissect the transcriptional basis of intergenerational plasticity. Our analyses identify regulators of high-temperature-induced dauer formation, maternal provisioning factors, and RNAi pathway components that together shape dauer responses across generations.

## Results

### Progeny of temperature-induced post-dauers exhibit increased dauer formation upon re-exposure

*C. elegans* nematodes are maintained at 20°C under laboratory conditions and enter diapause as dauer larvae when exposed to 27°C (*Ailion and Thomas*, 2000). We asked whether parental (P0) entry into dauer in response to elevated temperature influences the propensity of subsequent generations to enter dauer. We quantified dauer entry in the first (F1) and second (F2) generations of parental hermaphrodites (P0) exposed to 27°C (**Figure 1A**). Animals were exposed as embryos to 27°C for 45 hours, inducing dauer formation in the mean of 25.8% ± 1.9 SEM (**Figure 1B**). Following incubation, animals were separated into post-dauers (*PD-temp*; individuals that entered dauer at 27°C) and non-dauer (*ND-temp*; individuals that did not enter dauer at 27°C) groups, transferred to fresh plates, and allowed to develop to adulthood.

**Figure 1.**
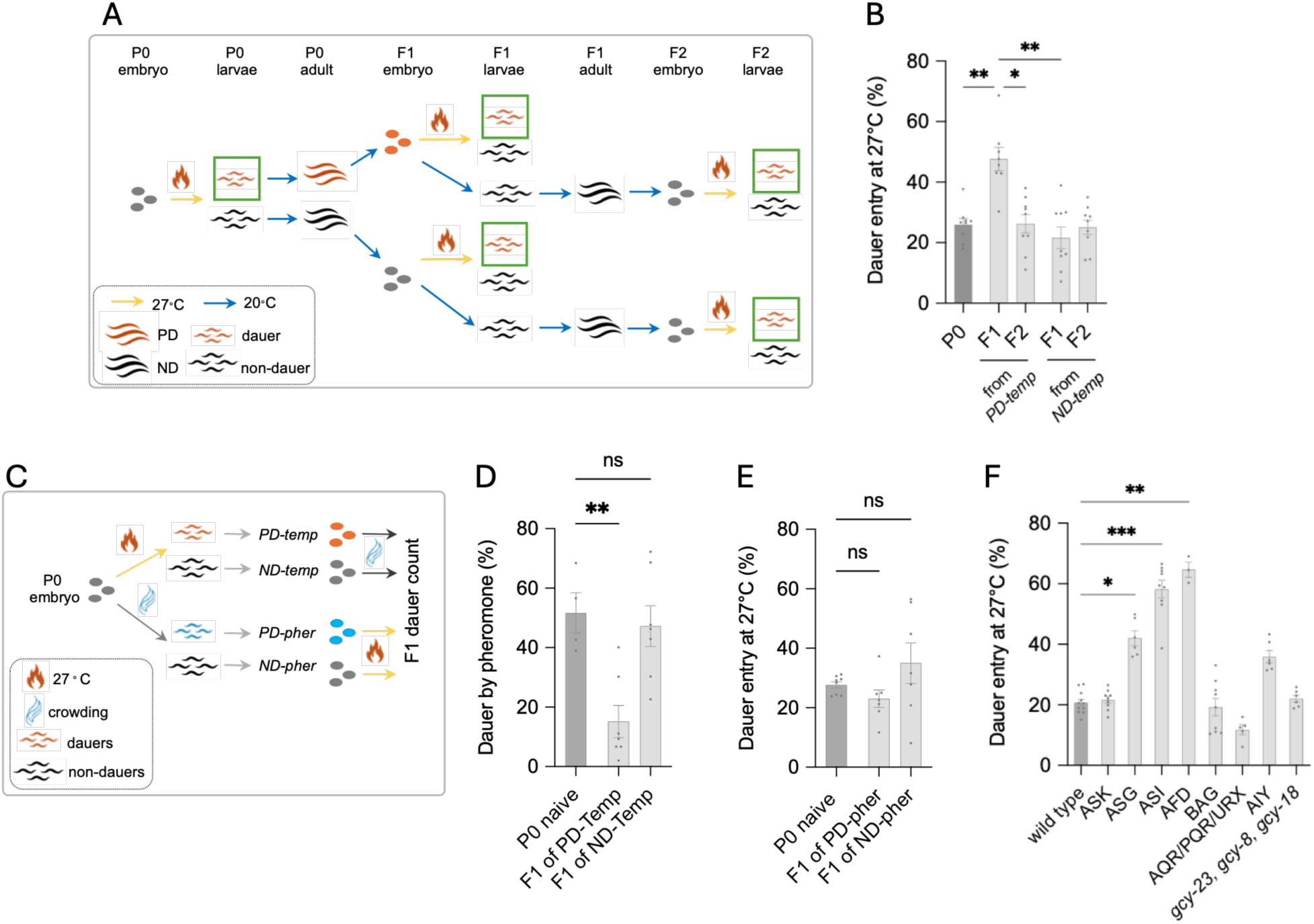
High-temperature diapause elicits a stimulus-specific intergenerational increase in dauer formation. (A) Experimental design to assess inter-and transgenerational consequences of embryonic heat exposure. Embryos (P0) were incubated at 27 °C for 45 h. Temperature-induced post-dauers (*PD-temp*) and non-dauers (*ND-temp*) were separated and propagated to generate F1 and F2 progeny. (N=9, 30-100 animals) (B) Percentage of dauer entry in naïve P0 animals, F1 progeny derived from *PD-temp* or *ND-temp* parents, and F2 progeny derived from the corresponding F1 lineages following re-exposure to heat (N = 9 biological replicates). Statistical significance was assessed using a Kruskal–Wallis test followed by Dunn’s multiple comparisons test (all pairwise comparisons); only significant differences are shown. Error bars represent mean ± SEM. (**C**) Experimental design to test the stimulus specificity of dauer induction in the F1 generation. P0 embryos were exposed either to heat (27 °C, 45 h) or to a pheromone mix. The resulting PD and ND animals were allowed to reach adulthood, and their progeny were subsequently challenged with the alternate stimulus. (**D–E**) Percentage of dauer entry induced by heat in naïve P0 animals and in F1 progeny derived from pheromone-exposed parents (*PD-crowd* and *ND-crowd*) (N = 7 biological replicates). Statistical significance was assessed using a Kruskal–Wallis test followed by Dunn’s multiple comparisons test (all pairwise comparisons); only significant differences are shown. Error bars represent mean ± SEM. (**F**) Percentage of dauer entry at 27 °C in wild-type animals and in animals with genetic ablations in specific neurons or mutations in thermosensory components (N > 3 biological replicates). Statistical significance was assessed using a Kruskal–Wallis test followed by Dunn’s multiple comparisons test (all comparisons versus wild type); only significant differences are shown. Error bars represent mean ± SEM.

Progeny of *PD-temp* parentals exhibited a significantly higher rate of dauer entry upon intergenerational re-exposure to 27°C, averaging 47.6% ± 3.8% SEM, indicating that parental dauer experience enhances dauer formation in the immediate generation. By contrast, progeny of *ND-temp* parentals displayed dauer formation with a mean of 21.6% ± 3.57 SEM (**Figure 1B**), similar to naive parental animals. In the F2 generation—whose F1 parents had not been exposed to high temperature—dauer formation for F2 from *PD-temp* P0s was 26.2% ± 3.0% SEM, while dauer formation for F2 progeny from *ND-temp* P0s was 25.1% ± 2.4% SEM (**Figure 1B**). Together, these results indicate that the increased propensity for dauer is intergenerational and reversible, requiring only a single stress-free generation for reset.

Because high temperature-induced diapause in P0s increased dauer penetrance in the F1 generation when F1 were exposed to high temperature, we asked whether dauer penetrance in the F1 of *PD-temp* P0s would similarly be increased when the F1s are exposed to other diapause-inducing cues—specifically pheromone. To test this question, we exposed the F1 progeny of *PD-temp* animals to pheromone-plates (2%v/v) in 25°C incubators for 48 hours (**Figure 1C**). Notably, only 15.2% ± 5.4% SEM, of F1 animals from *PD-temp* P0 entered dauer under pheromone exposure, whereas 51.7% ± 6.8% SEM, of F1 progeny from *ND-temp* P0 formed dauers—levels comparable to naïve animals that form dauers at a mean of 47.2% ± 6.9% SEM, (**Figure 1D**). Thus, parental experience with heat-induced diapause enhances dauer formation upon intergenerational heat exposure but suppresses dauer entry in response to pheromone.

To determine whether this asymmetry extends to pheromone exposure, we next examined progeny of animals subjected to pheromone-induced dauer conditions. Here, the F1 progeny of pheromone-induced post-dauer P0 formed dauers in a mean of 23.0% ± 2.9% SEM when exposed to high 27°C temperature, rates comparable to naive animals exposed to 27°C (mean of 27.7% ± 1.1% SEM) (**Figure 1E**). Similarly, the F1 progeny of pheromone-exposed non-dauer P0s exhibited a mean of 35% ± 6.8% SEM dauer formation when exposed to high 27°C temperature, with no difference in heat-induced dauer formation relative to naive animals exposed to heat (**Figure 1E**).

We also examined dauer formation under high-density conditions. Embryos were incubated at high population density (12,000 larvae with abundant *E. coli* OP50) at 25°C, and dauer formation was quantified after 70 hours. Following incubation, post-dauer animals (*PD-crowd*; individuals that entered dauer under high-density conditions) were then transferred to standard plates for recovery and progeny were re-exposed to high density conditions (**Figure S1A**). In the P0 generation, this treatment yielded 2.8% ± 0.7% SEM, dauers. Upon re-exposure to the same conditions, the F1 progeny of *PD-crowd* animals formed dauers at a similar frequency: 3.3 ± 0.53% SEM, indicating that, in contrast to thermal stress, parental exposure to high density conditions does not enhance dauer entry in progeny exposed to the same high-density conditions (**Figure S1B**). Together, these findings indicate that intergenerational effects on dauer formation are specific, with heat exposure—but not pheromone—eliciting a cue-specific modulation of progeny responses.

To assess whether the temperature input regulating HID is sensed by specific sensory neurons, we examined animals with genetic ablation of specific neurons and exposed them to 27°C. Loss of AFD produced the strongest increase in dauer formation, from 20.7% ± 1.1% SEM, in wild type to a mean of 64.6% ± 2.5% SEM (consistent with AFDs known roles in temperature sensation) (*Goodman and Sengupta*, 2018; *Clark et al.*, 2006), while loss of ASI and ASG also significantly elevated HID to 58.2% ± 2.9% SEM, and 42.1% ± 2.5% SEM, respectively (**Figure 1F**). By contrast, ablation of other tested sensory neurons, including ASK, BAG, and AQR/PQR/URX, as well as mutation of the AFD-associated thermotaxis-defective triple mutant of receptor guanylyl cyclases *gcy-23 gcy-8 gcy-18*, did not significantly alter dauer formation. No tested ablation reduced dauer formation below wild-type levels. These findings suggest that HID is not driven by a single obligate sensory input but is instead normally restrained by a subset of sensory neurons, with AFD exerting the strongest modulatory effect.

### Temperature-induced diapause induces distinct transcriptional states across generations

The stimulus-specific and transient intergenerational increase of dauers after heat exposure suggests that parental diapause experience is associated with transcriptional changes across generations. To uncover the molecular basis of this phenomenon, we performed RNA sequencing across three generations (P0– F2) and four experimental conditions: (1) crowding-induced post-dauer adults in the P0 generation (*P0-crowd*), (2) temperature-induced post-dauer adults in the P0 generation (*P0-temp*), (3) temperature-induced F1 post-dauer adults from *P0-temp* parents (*F1-temp*), and (4) temperature-induced F2 post-dauer adults descended from rested F1 that were not exposed to high temperature following *P0-temp* (*F2-temp*). *P0-crowd* was used as a comparative control to distinguish transcriptional changes associated with temperature-induced diapause from those associated with crowding-induced dauer formation.

Differential expression analysis (DEA) revealed a marked transcriptional divergence between parental post-dauer adults derived from temperature-induced diapause (*P0-temp*) and crowding-induced dauer formation (*P0-crowd*). Comparing *P0-temp* and *P0-crowd* animals identified 2,919 differentially expressed genes (Padj < 0.001; |log₂FC| ≥ 2), with 792 upregulated and 2,127 downregulated in *P0-temp* animals (**Dataset 1**). Principal component analysis (PCA) from gene expression values, clearly separated these groups (PC1=76.8%, PC2=10.1%), underscoring stimulus-specific molecular responses (**Figure 2A**). Similar separation was observed in the *F1-temp* (PC1=44.5%, PC2=30.3%; **Figure S2A**) and *F2-temp* (PC1=44.4%, PC2=17.7%; **Figure S2B**) comparisons (**Dataset 1**).

**Figure 2.**
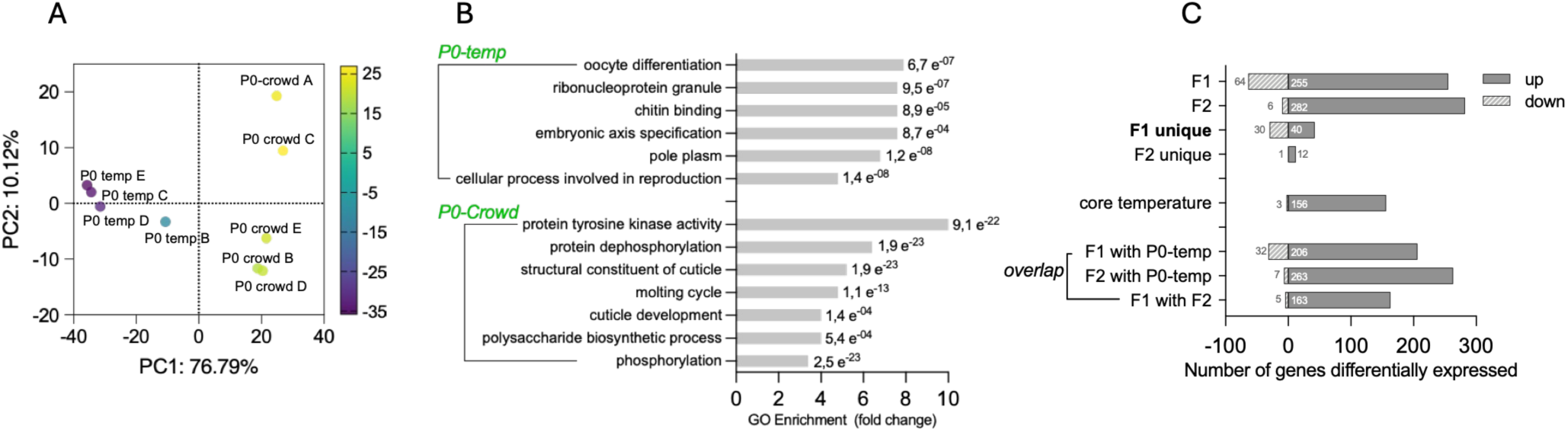
Temperature-induced diapause imprints a distinct and heritable adult transcriptional state. (**A**) Principal component analysis (PCA) of post-dauer (PD) adults reveals clear separation between temperature-induced (*PD-temp*) and pheromone-induced (*PD-crowd*) transcriptomes in the parental (P0) generation. The PCA was constructed using the 1,000 most variable genes. The proportion of variance explained is as follows: PC1, 76.79%; PC2, 10.12%. (**B**) Gene Ontology (GO) enrichment analysis highlighting stimulus-specific biological programs in *PD-temp* and *PD-crowd* adults. Q values were obtained using the WormBase Enrichment Analysis tool. Top hits were selected based on Q value. Full data is provided in Dataset 2. (**C**) Differentially expressed genes (DEGs) across P0, F1, and F2 post-dauer adults relative to the *P0-crowd* reference.

While adults from all samples were synchronously staged (see Methods), Gene Ontology enrichment analysis indicated that *P0-temp* adults were enriched for categories related to reproduction and oocyte differentiation while *P0-crowd* adults were enriched for signaling processes such as tyrosine kinase activity and protein dephosphorylation (**Figure 2B**). *P0-temp* adults were enriched for genes expressed in embryonic and oocyte lineages (**Figure S2C**) and associated with phenotypes such as abnormal P-granules or multiple nuclei in the early embryo. In contrast, genes upregulated in *P0-crowd* adults were enriched for expression in males and the hypodermis, linked to intestinal vacuolation and metabolic dauer traits (**Figure S2D**).

To contextualize our datasets, we integrated the P0–F2 transcriptomes with curated RNA-seq profiles of dauer versus reproductive development (*Lee et al.*, 2017), dauer entry induced by high temperature (*Corchado et al.*, 2026), and L2d, dauer, young adult, and adult stages from the *C. elegans* modENCODE project (*Gerstein et al.*, 2010). Principal component analysis comparing these datasets revealed that the P0, F1, and F2 transcriptomes did not fully overlap with canonical young adult profiles from the modENCODE dataset, despite all samples representing synchronized young adult animals (**Figure S2E**). Instead, our samples occupied an intermediate transcriptional space that partially overlapped with both young adult and dauer-associated developmental states. These observations suggest that prior developmental experience may leave a persistent transcriptional imprint that remains detectable in adulthood. Rather than reflecting a simple shift toward a dauer-like state, the transcriptomes of P0, F1, and F2 animals appear to retain features associated with multiple developmental trajectories, distinguishing them from adults that have not experienced dauer-inducing conditions.

We next examined the transcriptional landscapes of P0-F2 post-dauer adults derived from temperature-induced diapause. DEA relative to the P0 crowding-induced reference (*P0-crowd*) revealed distinct but overlapping transcriptional landscapes in *P0-temp*, *F1-temp*, *F2-temp*, and *P0-crowd* (**Figure S2F**). Specifically, we observed that the transcriptional profiles of *P0-temp* and *F1-temp* were most similar to each other, while *F2-temp* was most similar to *P0-crowd* by hierarchical clustering (**Figure S2F)**.

Hundreds of genes were differentially expressed in both *F1*-*temp* and *F2-temp* animals against *P0-crowd* (319 genes and 288 genes, respectively), including a shared “core” temperature-responsive set of 159 genes. Notably, we observed distinct generation-specific subsets: 70 regulated genes were unique to *F1-temp*, and 13 genes were unique to *F2-temp* (**Figure 2C, Dataset 2**). Together, these findings indicate that post-dauer adults across generations display extensive but partially overlapping transcriptional responses following high-temperature-induced diapause.

### F1-specific signatures reveal genes that promote dauer entry at high temperature

The broader transcriptional profile of post dauer animals is consistent with the possibility that developmental challenge remodels the adult physiological state, providing a potential explanation for the transient enhancement of dauer formation observed in the F1 generation. To identify genes that might contribute to this response, we compared differentially expressed genes across the P0, F1, and F2 generations, using *PD-crowd* animals as the reference condition. This analysis identified a set of 40 genes uniquely upregulated in the F1 generation (**Table 1**), potentially representing transient regulatory programs underlying the increased dauer formation observed in the progeny of heat-induced dauers. Indeed, of these F1-upregulated genes, nine overlap with transcripts previously associated with dauer formation (*Lee et al.*, 2017) (**Dataset 3**), and 19 correspond to genes highly expressed in dauer larvae according to developmental stage profiles from the modENCODE dataset (*Gerstein et al.*, 2010).

**Table 1.**
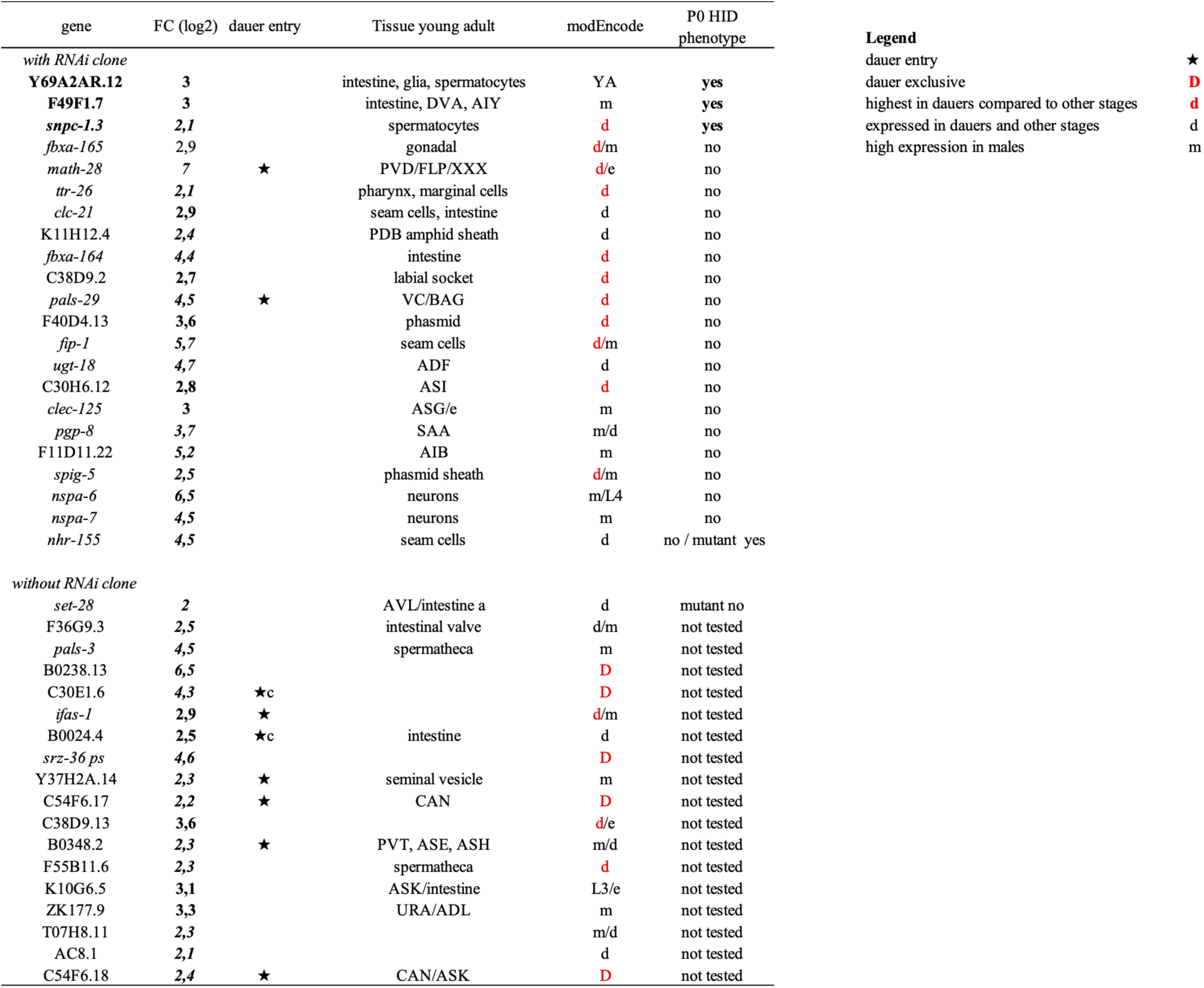
Transcripts uniquely upregulated in F1 post-dauer adults derived from high-temperature diapause (*PD-temp*). Columns indicate the gene name, fold change relative to *PD-crowd*, whether the gene has previously been reported to be required for dauer entry (*Lee et al.*, 2017), **Dataset 3**), the tissue in which the gene is predominantly expressed during the young adult stage in at least 5% of cells (*Ghaddar et al.*, 2023), the developmental stage where the expression is highest according to modENCODE data (*Gerstein et al.*, 2010), and the HID phenotype observed in this study. The table also indicates whether the RNAi clones used are predicted to target more than one gene.

For 20 of these candidate genes, RNAi clones were available for functional analysis (*Fraser et al.*, 2000; *Kamath et al.*, 2003). To assess the contribution of these F1-enriched transcripts to the intergenerational phenotype, animals were exposed to gene-specific RNAi for two consecutive generations, thereby ensuring gene knockdown during the F1 generation, when enhanced dauer formation is observed. An initial RNAi screen identified three genes whose silencing altered dauer formation at 27°C (**Figure S3**). Independent validation confirmed that knockdown of F49F1.7, Y69A2AR.12, or *snpc-1.3* significantly reduced dauer formation at 27°C (**Figure 3A**), supporting a role for these genes in promoting dauer entry under high-temperature conditions. To determine whether these genes specifically contribute to the transient increase in dauer formation observed in the F1 generation following heat-induced dauer, we next sought genetic validation. A loss-of-function allele was available only for *snpc-1.3*. Mutant animals formed dauers at frequencies comparable to wild type in the parental generation (P0; 27.3%), indicating that *snpc-1.3* is not generally required for temperature-induced dauer formation. In contrast, *snpc-1.3* mutants failed to exhibit the enhanced dauer formation characteristic of the F1 generation, producing only 4.5% dauers on average (**Figure 3B**). These findings identify *snpc-1.3* as a specific requirement for the transient intergenerational increase in dauer formation following heat exposure. Comparable genetic validation could not be performed for F49F1.7 or Y69A2AR.12, as mutant strains were not available.

**Figure 3.**
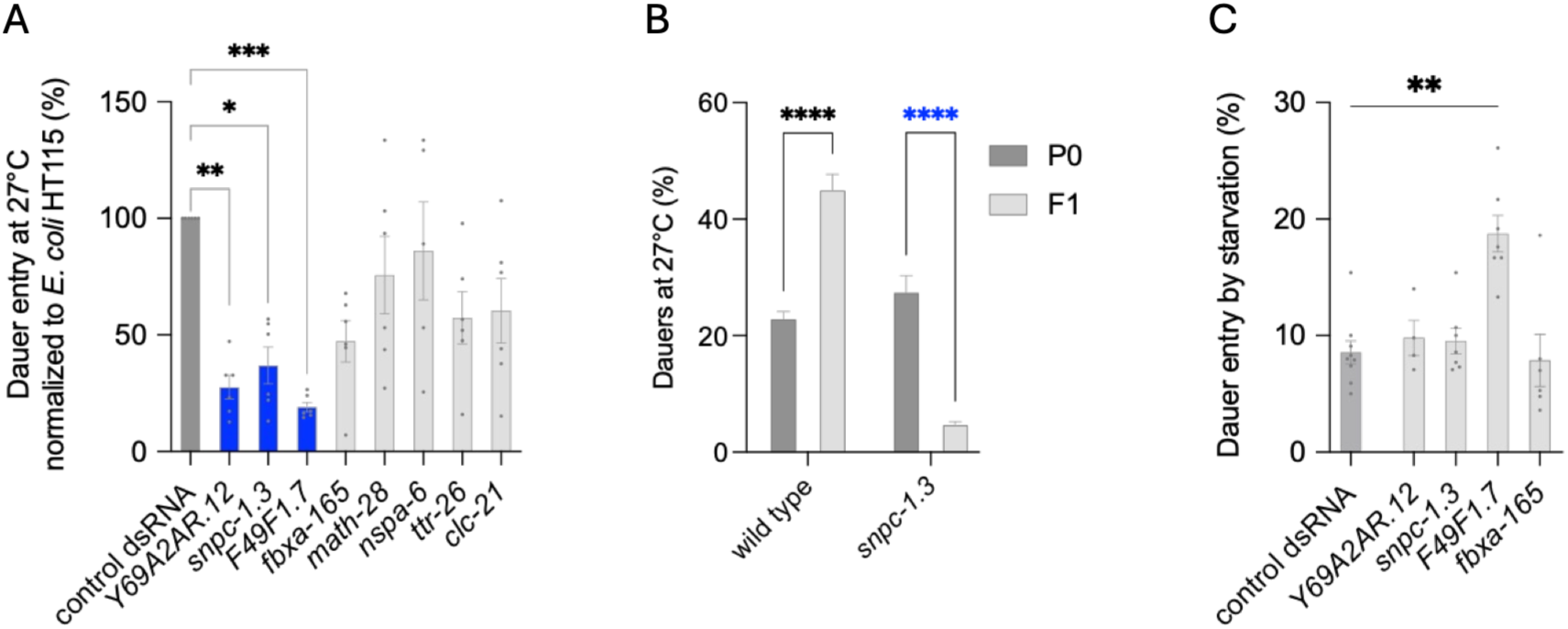
Identification and functional testing of F1-upregulated, heat-inducible dauer genes. (A) Dauer formation at 27 °C under RNAi treatment (N = 6 biological replicates). Statistical significance was assessed using a Kruskal–Wallis test followed by Dunnett’s multiple comparisons test (all comparisons versus control); only significant differences are shown. Error bars represent mean ± SEM. (B) Intergenerational HID assays of *snpc-1.3,* showing dauer entry in F1 progeny of heat-exposed parents. Statistical significance was assessed using an ordinary two-way ANOVA followed by uncorrected Fisher’s LSD, with a single pooled variance. Comparisons between F1 and P0 were performed. Error bars represent mean ± SEM. (N=7 or more biological replicates). (**C**) Dauer formation under starvation conditions (2 weeks) after RNAi treatment across three generations (N = 7 biological replicates). Statistical significance was assessed using the Kruskal–Wallis test followed by Dunn’s multiple comparisons test, with comparisons made against the control.

To determine whether these genes also regulate dauer formation under a distinct environmental challenge, RNAi-treated animals were subjected to starvation-induced dauer formation. Silencing of Y69A2AR.12 or *snpc-1.3* did not significantly alter starvation-induced dauer formation, whereas knockdown of F49F1.7 increased dauer formation under starvation conditions (**Figure 3C**). Together, these findings suggest that the functions of these genes are not universally required for dauer formation but instead differ according to the inducing stimulus. In particular, our results identify *snpc-1.3* as a specific regulator of the transient intergenerational increase in dauer formation following heat exposure. Whether F49F1.7 and Y69A2AR.12 also contribute to this F1-specific response will require genetic validation in future studies.

### Temperature-induced diapause represses *vit-3*, a negative regulator of dauer formation

In addition to genes induced by high-temperature diapause, we also examined transcripts that were consistently repressed in the F1 *PD-temp* generation. Comparative transcriptomics revealed a significant downregulation of all vitellogenin genes in F1 *PD-temp* compared to P0 *PD-temp* (**Table S1**). Notably, *vit-3* is the only transcript that in the F1 *PD-temp* generation that is downregulated relative to both P0 *PD-temp* (FC = -3.9, Padj = 2.09E-09) and F2 *PD-temp* (FC = -2.9, Padj = 0.0048). This pattern was also evident when comparing normalized counts of *P0-temp*, *F1-temp*, and *F2-temp* against *PD-crowd* controls (**Figure 4A**). *vit-3*, *vit-4*, and *vit-5* encode the yolk protein Y170A (*Blumenthal et al.*, 1984). Altered yolk provisioning has previously been linked to intergenerational effects on developmental timing, starvation responses, and other inherited physiological traits (*Perez and Lehner*, 2019). These observations prompted us to test whether vitellogenins modulate dauer formation at elevated temperature in the parental generation. *vit-3* mutants formed significantly more dauers at 27°C (**Figure 4B**) and under starvation (**Figure 4C**) indicating that *vit-3* normally inhibits dauer entry under thermal stress and starvation. By contrast, *vit-2 vit-1* double mutants disrupting the Y170B (*Blumenthal et al.*, 1984) protein did not show altered dauer formation, suggesting gene-specific roles among the vitellogenins (**Figure 4C**). We next validated the transcriptional changes in the F1 compared to the P0 generation using a fluorescent reporter with mCherry knocked into the C-*terminus* of *vit-3* (*Zhai et al.*, 2022) in animals exposed to 27°C. Reporter quantification across developmental time points—including P0 and F1 embryos, L2 larvae, and dauer larvae—showed that *vit-3* expression is reduced in F1 embryos and L2 descended from *P0-temp* parentals compared to their *P0-temp* parentals, while dauers display similar levels in both generations as P0 and F1 (**Figure 4D, 4E**). These results indicate that maternal VIT-3 signaling negatively regulates dauer entry in progeny of temperature induced dauer animals exposed to high temperature.

**Figure 4.**
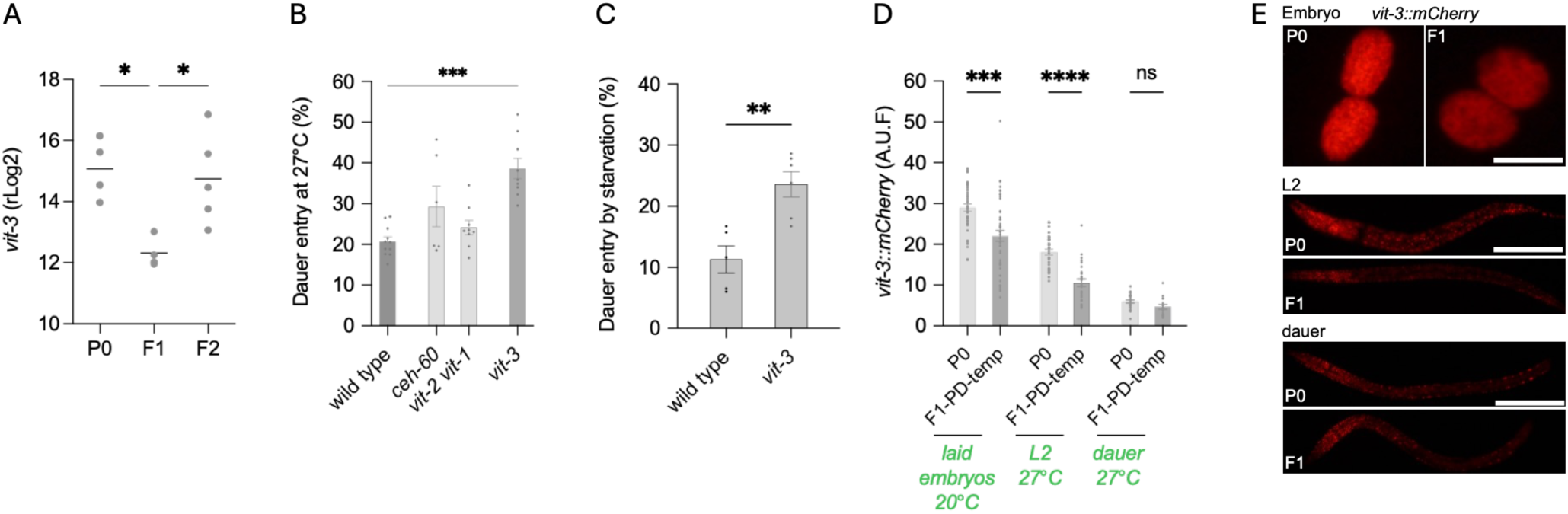
Temperature-induced diapause represses *vit-3* and promotes dauer formation in F1 progeny. (**A**) Differential expression analysis of *vit-3* across P0, F1, and F2 generations of temperature-induced post-dauer animals. Regularized log₂ (rlog) values were obtained using DESeq2 (N = 4 biological replicates for P0 and F1; N = 5 for F2). Statistical significance was assessed using a Kruskal-Wallis test followed by Dunn’s multiple comparisons test (all pairwise comparisons); only significant differences are shown. Error bars represent mean ± SEM. (**B**) Dauer formation at 27°C in *vit-3* mutants and *vit-2 vit-1* double mutants (N = 9 biological replicates). Statistical significance was assessed using a Kruskal– Wallis test followed by Dunnett’s multiple comparisons test (all comparisons versus control); only significant differences are shown. Error bars represent mean ± SEM. (**C**) Starvation-induced diapause in *vit-3* animals compared to control animals. N= 6 biological replicates. Statistical significance was assessed using a nonparametric Mann Whitney test. (**D**) mCherry fluorescent protein fusion to C-*terminus* of VIT-3 reporter expression across parental and F1 generations. At least N = 20 animals per condition were scored. Statistical significance was assessed using Welch’s one-way ANOVA (Brown– Forsythe) followed by Dunnett’s T3 multiple comparisons test. Comparisons between F1 and P0 were performed within the same developmental stage. Error bars represent mean ± SEM. (**E**) Representative images of protein fusion reporter of *vit-3* animals across parental and F1 generations in embryos, L2 and dauers. The scale bar is 50 µm.

### RNAi pathways coordinate thermal diapause in the parental generation and mediate intergenerational priming

Adaptive developmental responses to stress frequently rely on small-RNA pathways acting cell-autonomously, systemically, and across generations (*Bharadwaj and Hall*, 2017; Palominos et al., 2017; *Serey et al.*, 2025). To determine whether small-RNA-mediated regulatory mechanisms contribute to high-temperature-induced diapause, we first examined dauer formation in response to heat exposure in mutants affecting distinct functional modules of small-RNA biology. These included factors required for cell-autonomous and systemic RNAi, nuclear RNAi and inherited silencing, RNA-dependent siRNA amplification, and the biogenesis or function of specific small-RNA classes, including piRNAs, spermatogenesis-associated 26G-RNAs, and germline 22G-RNAs. Specifically, we analyzed mutants in *rde-1* and *rde-4* (cell-autonomous RNAi), *sid-1* and *sid-2* (systemic RNAi), *nrde-2* and *nrde-3* (nuclear RNAi), *rrf-1* and *mut-16* (siRNA amplification and processing), *hrde-1* and *znfx-1* (heritable small-RNA pathways), *prg-1* (piRNA-mediated regulation), *alg-3*; *alg-4* (spermatogenesis-associated 26G-RNA biogenesis and function), and *wago-1* (germline 22G-RNA-mediated silencing). We also examined *lin-15b* mutants, which display enhanced RNAi sensitivity. We also included the histone methyltransferases SET-18 and SET-25 because of their established roles in pathogen-induced dauer formation, where SET-18 contributes to response initiation and SET-25 to the maintenance of intergenerational memory (*Legüe et al.*, 2022).

All mutants yielded at least 400 live animals after heat treatment (**Figure S5A**). *mut-16* mutants showed impaired temperature-induced dauer formation, as previously reported (*Bharadwaj and Hall*, 2017). Interestingly, the two *rde-1* alleles produced distinct phenotypes. The *rde-1(ne219)* allele, a missense mutation affecting a conserved residue within the PAZ domain, impaired HID, whereas *rde-1(ne300)*, which introduces a premature stop codon within the PIWI domain, did not. Thus, disruption of RDE-1 does not uniformly impair thermal diapause, revealing an allele-specific requirement that may reflect distinct effects of the two mutations on RDE-1 function. *znfx-1* mutants also showed reduced dauer formation, suggesting that mechanisms linked to small-RNA inheritance contribute to HID. Nuclear RNAi components diverged: *nrde-2* mutants displayed a significant increase in dauer formation, whereas *nrde-3* mutants were indistinguishable from wild type. The *alg-3; alg-4* double mutant, as well as *wago-1* and *lin-15b* mutants, also displayed increased dauer formation (**Figure 5A**). *set-18* and *set-25* mutants were indistinguishable from wild type under these conditions. In contrast to HID, starvation-induced diapause was not affected in *rde-1(ne219)* or *znfx-1* mutants (**Figure S5B**). We next asked whether small-RNA pathways contribute to the enhanced F1 response to thermal stress. Because some mutants exhibited severe fertility and post-dauer recovery defects after heat-induced diapause, only a subset of genotypes could be readily propagated and tested in the next generation (*nrde-2*, *nrde-3*, and *hrde-1*). Consecutive HID assays showed that, unlike wild-type animals, *nrde-2* mutants failed to exhibit enhanced dauer formation in the F1 generation (**Figure 5B**). This indicates that NRDE-2 is required for intergenerational priming of dauer entry unlike NRDE-3 and HRDE-1 whose mutants retained F1 priming.

**Figure 5.**
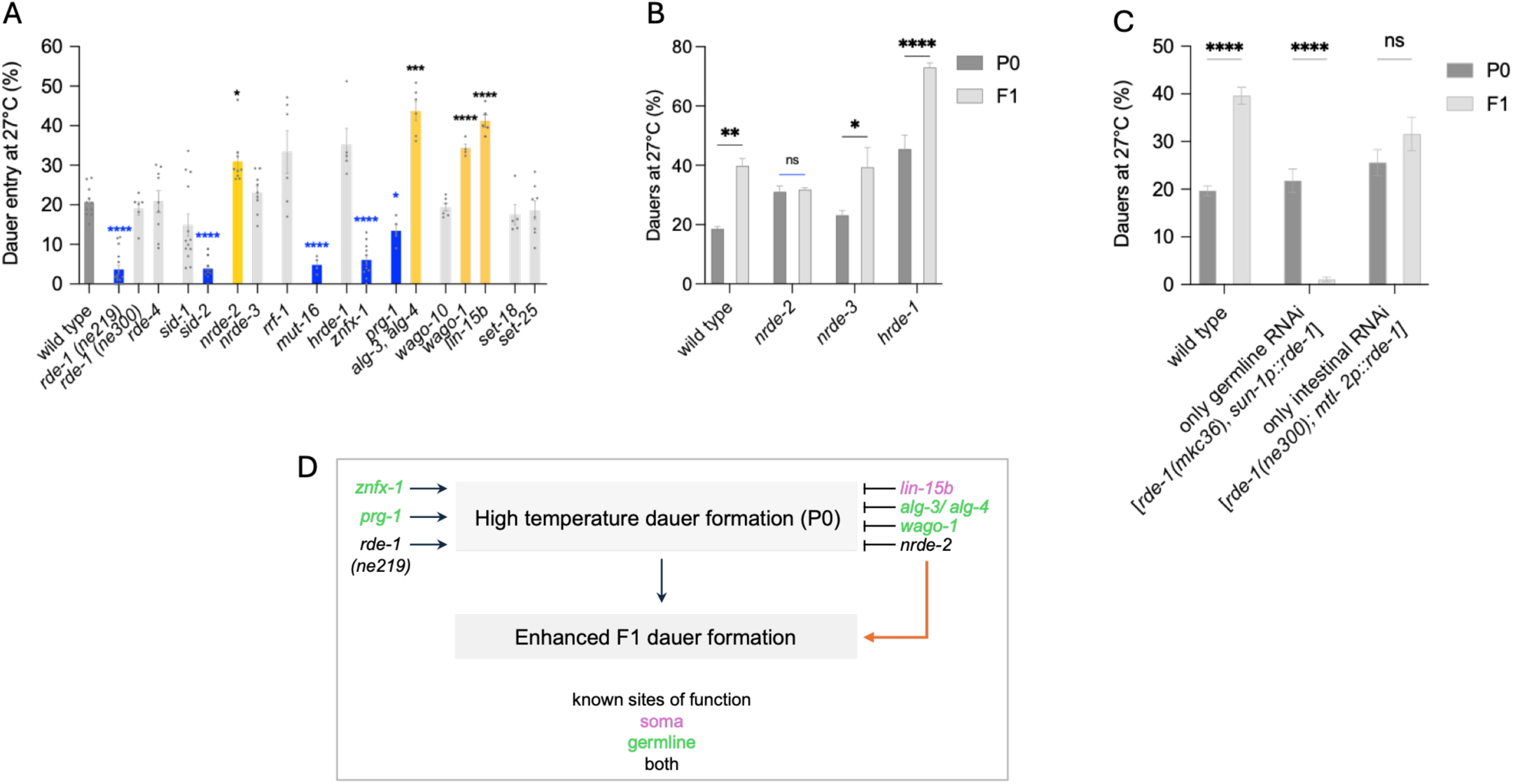
RNAi pathways coordinate thermal diapause in parental animals. **(A)** Dauer formation at 27 °C in mutants defective in different steps of the RNAi pathway (N = 4 biological replicates). Statistical significance was assessed using Welch’s one-way ANOVA (Brown– Forsythe) followed by Dunnett’s T3 multiple comparisons test (all comparisons versus control); only significant differences are shown. Error bars represent mean ± SEM. (**B**) Intergenerational HID assays of *nrde-2, nrde-3, hrde-1* showing dauer entry in F1 progeny of heat-exposed parents (N = 3 for F1 and N = 9 for P0). Statistical significance was assessed using an ordinary two-way ANOVA followed by Šídák’s multiple comparisons test. Comparisons between F1 and P0 were performed; only significant differences are shown. Error bars represent mean ± SEM. (**C**) Intergenerational HID assay using the tissue-specific expression of *rde-1* in the germline (using the *sun-1* promoter) and intestine (using the *mtl-2* promoter) (N = 4 for F1 and N = 7 for P0). Statistical significance was assessed using an ordinary two-way ANOVA followed by Šídák’s multiple comparisons test. Comparisons between F1 and P0 were performed; only significant differences are shown. Error bars represent mean ± SEM.

Having established that small-RNA pathways contribute to intergenerational priming and identified NRDE-2 as a key requirement, we next asked whether RNAi activity in specific tissues was sufficient to support this response. To address this question, we used strains in which RNAi activity was restricted to either the germline or the intestine. For the germline, we used a rescue strain generated in the RNAi-deficient *rde-1(mkc36)* null background (*Zou et al.*, 2019), in which RNAi function is restored exclusively in germ cells. For the intestine, we used a strain expressing *rde-1* under the intestinal *mtl-2* promoter. We exposed embryos to 27°C, both P0 and the F1 progeny of P0 *PD-temp* and quantified their ability to diapause in both generations. While either rescue was sufficient to support dauer formation in P0 animals, active RNAi in the germline alone failed to support enhanced dauer formation in the F1 progeny, and intestinal RNAi alone was similarly insufficient to prime descendants for increased dauer formation in the F1 (**Figure 5C**).

Together, these findings identify small-RNA pathways as key regulators of thermal diapause and its intergenerational enhancement. Multiple small-RNA pathway components modulate dauer formation in heat-exposed parental animals, whereas intergenerational priming in the F1 generation specifically requires the nuclear RNAi factor NRDE-2. Furthermore, restricting RNAi activity to either the germline or intestine alone was insufficient to support enhanced dauer formation in descendants, indicating that transmission of thermal stress information depends on coordinated RNA-based signaling across tissues.

### Diapause preserves reproductive fitness under heat and crowding stress

During the *C. elegans* diapause, germline stem cells arrest the cell cycle completely, thereby protecting reproductive capacity until conditions improve (*Narbonne and Roy*, 2006; *Kadekar and Roy*, 2019). When exposed to heat, pre-reproductive larvae and their descendants exhibit a marked decline in fertility, both in natural strains and in the domesticated N2 laboratory strain (*Petrella*, 2014). We hypothesized that diapause triggered by either high temperature or crowding could protect fertility once animals resumed development. We induced dauers by exposing embryos to high temperature (27°C, (*Ailion and Thomas*, 2000) or crowding (12,000 larvae at 25°C). After the perturbations, we separated dauers from non-dauers and each individual animal was then transferred to 20°C and allowed to develop to adulthood. Brood size was quantified over the first four days of fertility in all conditions.

Animals that did not enter dauers after exposure to heat (*ND-temp*) or pheromone (*ND-crowd*) produced significantly fewer progeny than controls raised continuously at 20°C (mean brood size = 104), indicating that early stress impairs reproductive capacity (**Figure 6A**). However, post-dauers from the same treatments (*PD-temp* and *PD-crowd*), although producing smaller broods than non-stressed controls, showed markedly larger broods than their non-dauer siblings (**Figure 6A**), indicating that dauer transit helps preserve fertility. We next quantified sterility, defined as adults without viable progeny within four days after reaching adulthood. *ND-temp* animals exhibited high sterility (46%), whereas only 8% of their *PD-temp* siblings were sterile (**Figure 6B**). A similar trend was observed following pheromone-induced dauer formation, with sterility reduced from 23% in *ND-crowd* animals to 2% in *PD-crowd* animals (**Figure 6B**). Together, these results indicate that diapause, regardless of the inducing stimulus, safeguards reproductive fitness.

**Figure 6.**
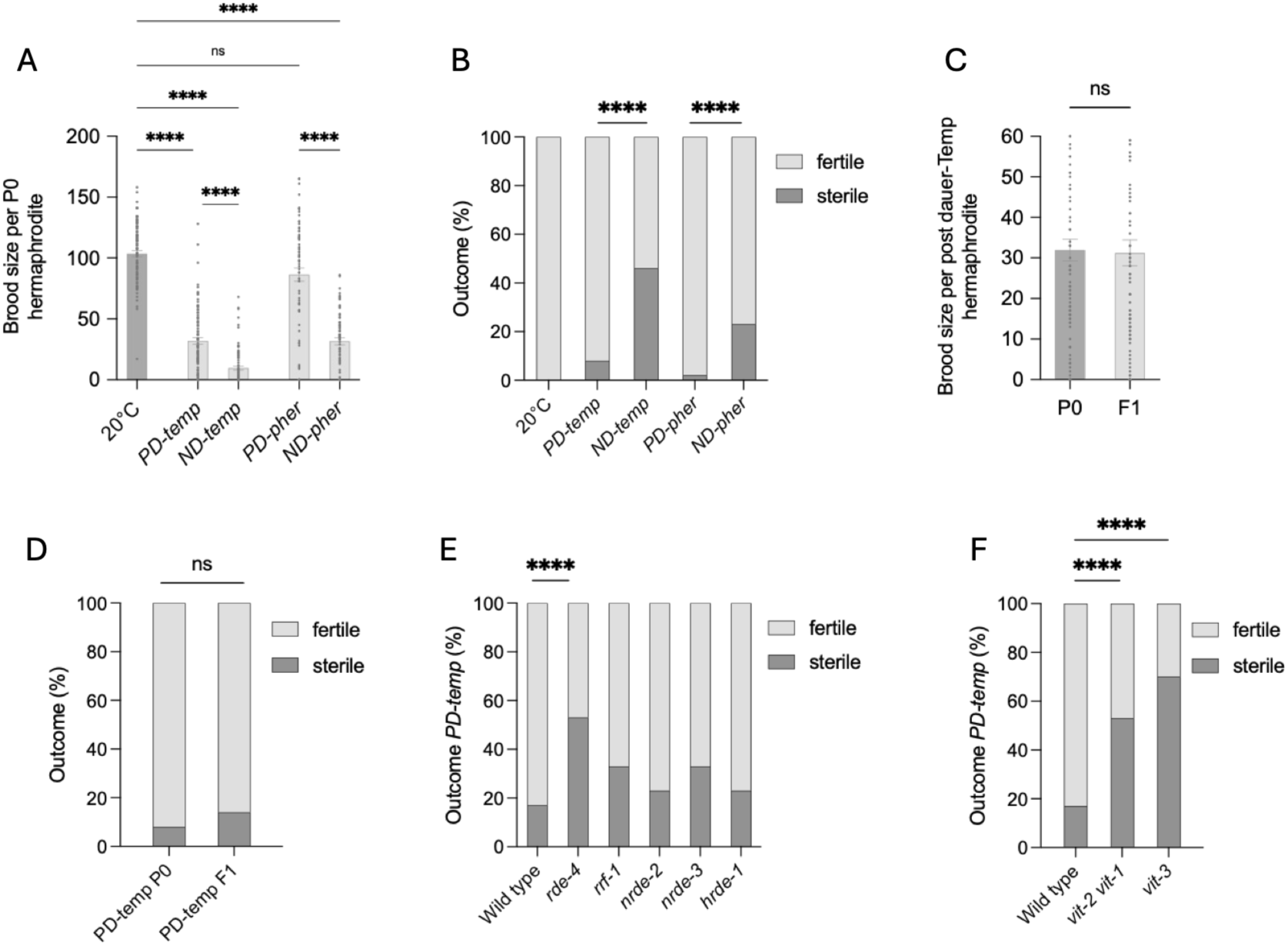
Diapause protects reproductive fitness under temperature and crowding. (**A**) Brood size of control animals grown at 20 °C and of post-dauer (PD) versus non-dauer (ND) animals that experienced high temperature (*PD-temp* and *ND-temp*; N = 90 each) or crowding (*PD-crowd* and *ND-crowd*; N = 70 each). Statistical significance was assessed using a Kruskal–Wallis test followed by Dunn’s multiple comparisons test. Error bars represent mean ± SEM. (**B**) Percentage of sterile adults under the same conditions (N = 90 for temperature and N = 70 for crowding, across three biological replicates). Statistical significance was assessed using Fisher’s exact test on cumulative counts from N = 3 biological replicates. (**C**) Brood size of *PD-temp* animals and their F1 progeny following re-exposure to high temperature (F1 *PD-temp*; N = 90 individuals). Statistical significance was assessed using a Mann–Whitney U test. Error bars represent mean ± SEM. (**D**) Percentage of sterile adults in *PD-temp* parents (P0) and their F1 progeny. Statistical significance was assessed using Fisher’s exact test on cumulative counts from N = 3 biological replicates (90 total animals per condition). (**E**) Percentage of sterile animals in mutants of RNAi pathway components after dauer recovery at 27 °C in the P0 generation (PD animals; N = 30 animals). Statistical significance was assessed using Fisher’s exact test on cumulative counts from N = 3 biological replicates. (**F**) Percentage of sterile animals in *vit-3* and *vit-2; vit-1* double mutants after dauer recovery at 27 °C in the P0 generation (PD animals; N = 30 animals). Statistical significance was assessed using Fisher’s exact test on cumulative counts from N = 3 biological replicates.

We next asked whether reproductive fitness was also preserved in the F1 of *PD-temp* parentals. F1 post-dauers showed brood sizes and sterility rates comparable to those of their P0 parents (**Figure 6C–D**), indicating that the reproductive protection associated with diapause is maintained following intergenerational heat exposure.

## Supplementary figures

**Figure S1.**
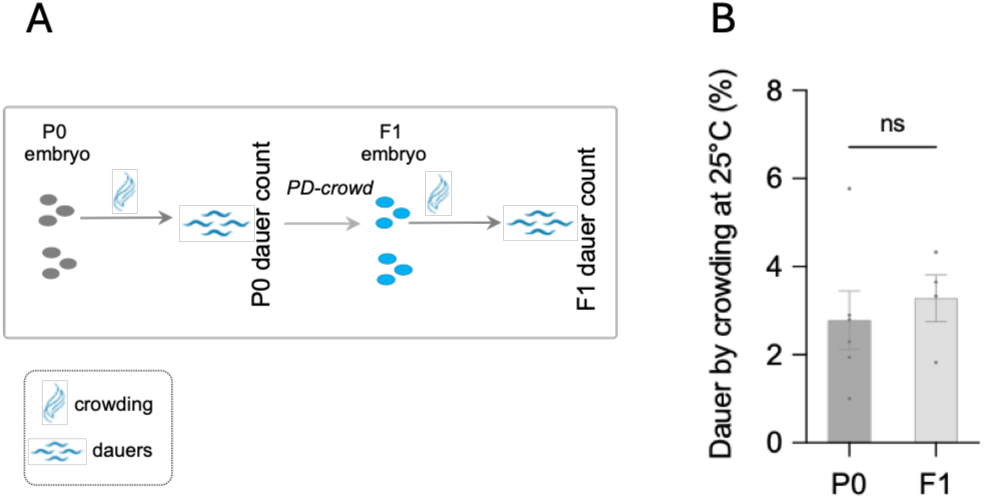
Percentage of dauer entry induced by high-density conditions. (A) Schematic diagram of the experimental design used to assess crowding treatments in naïve parental P0 animals and F1 progeny of *PD-crowd* animals. Crowding conditions consisted of high-density plates containing 12,000 larvae cultured with abundant *E. coli* OP50 at 25 °C. N=6 (B) 12,000 larvae with abundant *E. coli* OP50 at 25°C, in naïve animals and in F1 progeny derived from *PD-crowd* parents (N = 6 biological replicates for P0 and N = 4 for F1). Statistical significance was assessed using Mann–Whitney test. Error bars represent mean ± SEM. N=4

**Figure S2.**
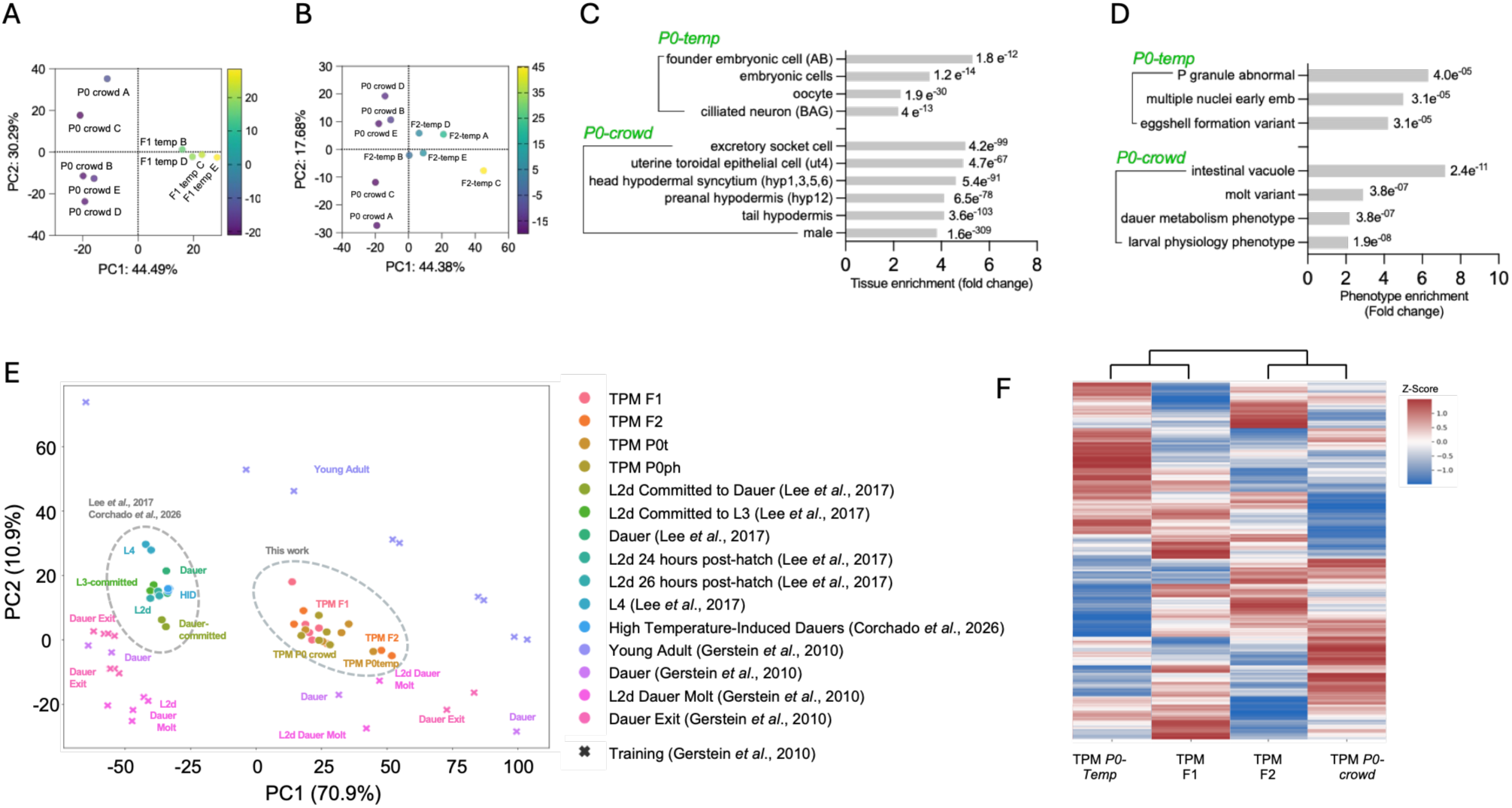
Transcriptomic profiling of *P0-temp*, *F1-temp*, and *F2-temp* relative to *P0-crowd*. (**A–B**) Principal component analysis (PCA) of the 1,000 most variable genes showing separation of post-dauer adults derived from *PD-temp* versus *PD-crowd* in the F1 (A) and F2 (B) generations. The proportion of variance explained is as follows: (A) PC1, 44.5%; PC2, 30.3%. (B) PC1, 44.4%; PC2, 17.7%. (**C**) Tissue enrichment analysis of genes upregulated in temperature-induced post-dauer adults (*PD-temp*, P0). Q values were obtained using the WormBase Enrichment Analysis tool. Top hits were selected based on Q value. Full data is provided in **Dataset 2**. (**D**) Phenotype enrichment analysis of genes upregulated in pheromone-induced post-dauer adults (*PD-crowd*, P0). Q values were obtained using the WormBase Enrichment Analysis tool. Top hits were selected based on Q value. Full data is provided in **Dataset 2**. (**E**) PCA plot of the variation in gene expression across the samples sequenced in this work, as well as publicly available RNA-seq datasets describing various *C. elegans* life stages (*Gerstein et al.*, 2010), dauer versus reproductive development (*Lee et al.*, 2017), and temperature-induced dauer development (*Corchado et al.*, 2026). The PCA was trained on the L2d dauer molt, dauer, post-dauer, L4, and young adult stages from (*Gerstein et al.*, 2010), using the variation across these stages as a reference. The training and remaining samples were then projected onto this PC space to investigate how the remaining samples vary along the principal component axes. The variations explained by PC1 and PC2 are listed in parentheses. Expression data (in tpm) were log-transformed, centered, and scaled before PC analysis. (**F**) Differential expression of 2932 genes across the conditions of this work. DEGs were selected from the P0 condition Padj < 0.001; |log₂FC| ≥ 2 of *P0-temp* vs *P0-crow*, Dataset 1. Expression data (in tpm) were log-transformed, centered, and scaled for differential expression analysis. Each row represents a single gene, and heat colors indicate expression level z-scores. Dendrogram indicates clustering of the conditions based on the similarity of their expression profiles, calculated using correlation distances and average-linked hierarchical clustering.

**Figure S3.**
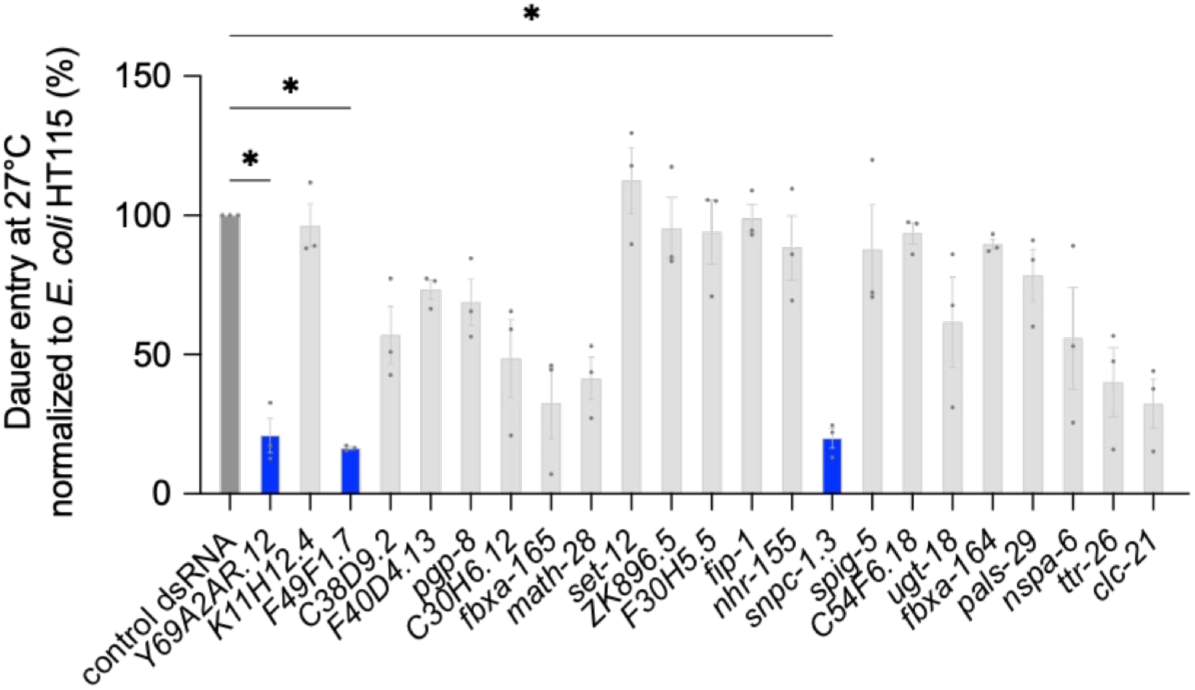
Screening of F1-specific genes and starvation-induced dauer formation. Dauer formation at 27 °C following RNAi treatment targeting 20 F1-upregulated candidate genes with available clones from the Ahringer library (*Fraser et al.*, 2000; *Kamath et al.*, 2003) (N = 3 biological replicates). Statistical significance was assessed using the Kruskal–Wallis test followed by Dunn’s multiple comparisons test, with comparisons made against the control.

**Figure S5.**
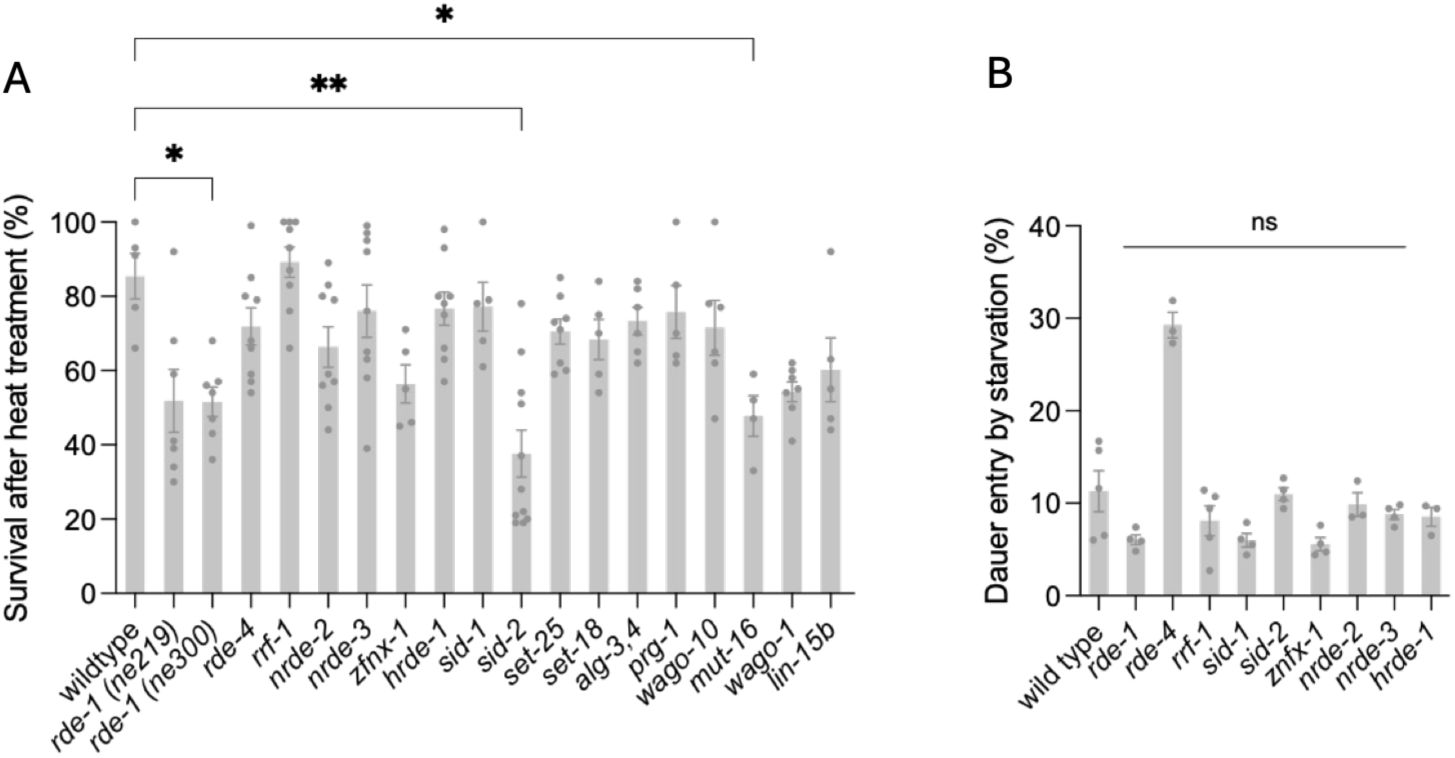
Survival and starvation responses of RNAi pathway mutants. (A) Survival following 27 °C–induced dauer formation in wild-type and mutant animals derived from bleached embryos incubated for 70 h at 27 °C. Survival was calculated as the proportion of animals alive relative to the initial number of eggs (N = 5 biological replicates). Statistical significance was assessed using the Kruskal–Wallis test followed by Dunnett’s multiple comparisons test (all comparisons versus control). (**B**) Dauer formation under starvation conditions. Strains were maintained under standard conditions and last fed 2 weeks prior to the starvation assay (N = 4 biological replicates). Statistical significance was assessed using the Kruskal–Wallis test followed by Dunnett’s multiple comparisons test (all comparisons versus control).

**Table S1.** Fold change of *PD-temp* across generations (*F1-temp* vs *P0-temp*, *F1-temp* vs *F2-temp* and *F2-temp* vs *P0-temp*) of vitellogenin genes.

| gene | *F1-temp* vs *P0-temp*<br>(Fold change) | Padj. | *F1-temp* vs *F2-temp*<br>(Fold change) | Padj. | *F2-temp* vs *P0-temp*<br>(Fold change) | Padj. |
| --- | --- | --- | --- | --- | --- | --- |
| *vit-1* | -3,2 | 3,08E-07 | 1,4 | 0,000204911 | ns | NA |
| *vit-2* | -2,5 | 3,24E-05 | ns | NA | ns | NA |
| ***vit-3*** | **-3,9** | **2,09E-09** | **-2,9** | **0,004804157** | **ns** | NA |
| *vit-4* | -3,8 | 1,27E-08 | 1,6 | 6,72E-05 | ns | NA |
| *vit-5* | -3,8 | 1,57E-10 | -1,1 | 0,002858001 | ns | NA |
| *vit-6* | -3,0 | 1,30E-06 | ns | NA | ns | NA |

## Datasets legends

**Dataset 1**. Differentially expressed genes in *P0-temp*, *F1-temp* and *F2-temp* compared to *P0-crowd*.

**Dataset 2**. Generation specific gene expression.

**Dataset 3**. List of genes differentially expressed in L2s vs dauers during dauer development at 25°C (PMID: 29167374).

**Dataset 4**. All data from the manuscript

**Dataset 5**. All statistics analysis in the manuscript

## Discussion

Our results identify HID as a distinct, stress-induced developmental program that couples dauer entry to two linked outcomes: preservation of reproductive capacity and a transient increase in dauer formation in progeny re-exposed to the same environmental condition. While preservation of reproductive fitness is shared with crowding-induced diapause, the intergenerational enhancement of dauer formation is transient and specific to thermal stress, distinguishing HID from chronic or long-term forms of environmental inheritance. Mechanistically, the intergenerational component depends on endogenous RNAi pathway activity and is associated with altered *vit-3* expression, although the relationship between these processes remains unresolved. Together, these findings indicate that the physiological and intergenerational consequences of dauer entry depend on the environmental cue that induces diapause.

High temperature is a strong developmental stress for *C. elegans*. While laboratory culture is typically maintained between 15° and 25°C (*Brenner*, 1974), natural substrates in nature can reach 4°C to 26°C (*Cook et al.*, 2017; *Crombie et al.*, 2019; *Crombie et al.*, 2022; *Crombie et al.*, 2024). Exposure to 26-27°C abruptly drops fertility in natural *Caenorhabditis* strains (*Petrella*, 2014), making dauer entry at 27°C a plausible strategy for preserving reproductive fertility. In this context, our results support a model in which thermal stress is first detected by sensory neurons, including AFD, ASI and ASG. Sensory perception is a key determinant of dauer entry (*Bargmann and Horvitz*, 1991), and temperature is a potent inducer of this developmental decision (*Golden and Riddle*, 1984). Although AFD neurons are the primary thermosensors in *C. elegans* (*Kimura et al.*, 2004), their specific role in high-temperature-induced dauer formation has remained unclear. Our results show that AFD ablation increases dauer formation under thermal stress, suggesting that AFD activity acts to restrain inappropriate developmental arrest. These observations suggest that sensory detection of elevated temperature is subsequently integrated with intracellular regulatory programs that determine whether animals commit to dauer entry.

Our RNAi screen identified several additional genes that may connect temperature sensing with dauer entry. Y69A2AR.12 encodes a predicted membrane-associated protein enriched in AFD and is negatively regulated by SKN-1 under basal conditions (Oliveira et al., 2009; Harris et al., 2023). F49F1.7 encodes a predicted secreted ShKT-domain protein associated with innate immune programs whose expression is induced and remains elevated following heat exposure (O’Rourke et al., 2006; Zhou et al., 2019). SNPC-1.3 is a male germline-enriched, SNAPc-related transcription factor required for spermatogenic piRNA expression and male fertility (Choi et al., 2021). Together with their RNAi phenotypes, these observations implicate genes associated with thermosensation, stress responses, and germline regulation in HID, although their tissues of action and underlying mechanisms remain to be determined. Beyond these candidates, HID also engages small-RNA pathways that shape parental dauer entry. *znfx-1* mutants showed reduced HID, implicating mechanisms associated with endogenous small-RNA regulation in dauer formation under thermal stress. *rde-1*, however, displayed a striking allele-specific phenotype. The *ne219* allele, which introduces an E414K substitution in RDE-1, impaired HID, whereas *ne300*, a strong loss-of-function allele carrying a premature stop codon (*Tabara et al.*, 1999), did not.

The allele-specific behavior of *rde-1* raises the possibility that the *ne219* phenotype reflects an altered RDE-1 activity rather than simple loss of function. Recent work suggests that the *ne219* mutant protein may retain the ability to interact with endogenous small RNAs, raising the possibility that the E414K substitution alters the repertoire or downstream activity of RDE-1-associated small RNAs in a manner relevant to thermal diapause (*Knittel et al.*, 2024). Interestingly, *ne219* also differs from *ne300* and a *rde-1* deletion in its effects on fertility at elevated temperature (*Knittel et al.*, 2024), suggesting that the functional consequences of this allele may become particularly relevant under thermal stress. At the same time, the increased dauer formation observed in mutants such as *wago-1*, *alg-3/alg-4*, *lin-15B*, and *nrde-2* suggests that multiple small-RNA pathways normally act to restrain dauer entry under thermal stress. Together, these findings support a model in which RNAi-related mechanisms tune the developmental threshold for dauer entry at elevated temperature. This interpretation is consistent with previous work showing that ALG-3/ALG-4-dependent 26G-RNAs and WAGO-associated 22G-RNAs contribute to the robustness of gene-expression programs under elevated temperature conditions (*Conine et al.*, 2010), (*Seroussi et al.*, 2023). Consistent with this idea, tissue-specific rescue experiments showed that RNAi activity restricted to either the germline or intestine was sufficient to support HID in the parental generation, whereas neither tissue alone was sufficient to support enhanced dauer formation in the F1 progeny. These findings point to a broader coordination of RNA-based regulation across tissues in the intergenerational response.

A subset of these small-RNA regulatory mechanisms extends into the intergenerational response. Among the factors tested, *nrde-2* plays a selective role in the enhanced dauer phenotype, distinguishing it from other nuclear RNAi components. This specificity, together with the absence of effects from histone methyltransferases such as *set-18* and *set-25*, previously implicated in inter-and transgenerational responses to pathogenic stress (*Legüe et al.*, 2022), suggests that the intergenerational increase in HID is more consistent with a short-term, reversible form of inheritance that depends on small RNA pathways without any suggestion of stable epigenetic reprogramming. This interpretation is further supported by the transient nature of the phenotype, which resets after a single stress-free generation, distinguishing this process from the long-term inheritance effects associated with chronic high-temperature exposure (*Klosin et al.*, 2017; *Frejacques et al.*, 2026).

In parallel to RNAi-dependent regulation, maternal provisioning by vitellogenins (*Perez and Lehner*, 2019) could contribute to the intergenerational response (*Perez et al.*, 2017; *Eroglu et al.*, 2024). While vitellogenins are broadly required for yolk formation (*Blumenthal et al.*, 1984), the specific involvement of *vit-3* in HID, but not *vit-1*, *vit-2*, or *ceh-60,* argues against a general effect of yolk provisioning. Instead, *vit-3* may act more selectively, potentially as a signaling factor that biases progeny toward dauer entry under thermal stress. Although previous studies have reported associations between NRDE-2 and VIT-3 (*Wan et al.*, 2020), our data do not resolve whether these factors act in a shared pathway. The most parsimonious interpretation is that RNAi dependent regulation and maternal provisioning represent parallel inputs into the intergenerational phenotype.

At the level of organismal outcomes, the molecular and cellular processes mentioned above give rise to two separable but coordinated consequences of HID. First, dauer entry protects reproductive capacity in the parental generation, buffering the detrimental effects of elevated temperature on fertility. This protective effect is not unique to thermal stress, as it is also observed following crowding-induced dauer, suggesting that preservation of reproductive potential represents a general property of diapause. In contrast, the intergenerational increase in dauer formation is specific to thermal stress and is transient across generations, indicating that not all consequences of dauer are shared across inducing cues. The increased propensity of progeny to enter dauer upon re-exposure to heat may therefore represent a reversible adaptive strategy that transiently lowers the threshold for developmental arrest under recurring environmental stress. More broadly, these findings support the idea that dauer is not a uniform developmental endpoint, but rather a context-dependent physiological state whose downstream consequences are shaped by the environmental conditions that induce it.

## STAR Methods

### KRT table

#### Resource availability

##### Lead contact

Further information and requests for resources and reagents should be directed to and will be fulfilled by the lead contacts Andrea Calixto and Esteban Retamales.

### Materials availability

This study did not generate new unique reagents.

### Experimental model and subject details

#### C. elegans

Wild type (N2 Bristol) hermaphrodite nematodes and mutant strains PS8438 [*syIs600* (*col-183p*::mCherry + *odr-1p*::GFP)], YY186 *nrde-2*(*gg91*) II, YY158 *nrde-3(gg66*) X, YY538 *hrde-1*(*tm1200*) III, YY996 *znfx-1*(*gg561*) II, RB1815 *vit-3* (*ok2348*)X, MQD2884 *vit-2*(*ok3211*) *vit-1*(*hq532*) X, MQD2775 [*vit-3* (*hq485*[*vit-3::mCherry*]) *vit-2* (*crg9070*[*<u>vit-2::gfp</u>*]) X], WM27 (*rde-1* (*ne219*) V),WM45 (*rde-1*(*ne300)* V; WM49 *rde-4* (*ne301*) III, DCL569 [*rde-1(mkcSi13)* [*sun-1p*::*rde-1::sun-1* 3’UTR + *unc-119*(+)] II; IG1839 [*frSi17* II; frIs7 IV; *rde-1* (*ne300*) V], NL2098 rrf-1(*pk1417*) I, HC271 [ccIs4251 I; *qtIs3 sid-2(qt42)* III; *mIs11* IV], NL3321*sid-1* (*pk3321*) V, VC584 *ceh-60* (*gk280*) X, MT17463 *set-25*(*ne5021*) III, CZ25708 *prg-1*(*ju1574*) I, JMC192 *wago-10* (*tor133*) V, VC767 *set-18* (*gk334*) I, NL1810 *mut-16* (*pk700*) I, RB1096 *wago-1*(*ok1074*) I, MT2495 *lin-15b* (*n744*) X, PY7505 oyIs84 [*gpa-4p*::TU#813 + *gcy-27p*::TU#814 + *gcy-27p*::GFP + *unc-122p*::DsRed], PS6025 *qrIs2* [*sra-9::mCasp1*], XA2262 *qaIs2241* [*gcy-36p::egl-1* + *gcy-35p*::GFP], RJP56 *rpIs3* [*gcy-33p*::GFP *vsIs33 dop-3*::RFP] V, JPS271 *vxEx265* [*gcy-8p*::ICE + *myo-2p:mCherry]*, JN579 *peIs579* [*ttx-3p::casp1* + *ttx-3p:*:Venus + *lin-44p*::GFP], JN2113 *peIs2113* [*gcy-21p*::mCaspase + *tax-4p*::NLS::YC2.60 + *lin-44p*::GFP], IK597 *gcy-23*(*nj37*) *gcy-8*(*oy44*) *gcy-18*(*nj38*) IV, were grown at 20°C and *E. coli* OP50 (*Brenner*, 1974) in HERAtherm Thermo Scientific (IMC18) incubators, for at least 3 generations before experimentation with different temperatures. We used synchronized populations of nematodes to start the experiments and for the scoring. Details are provided below for each case.

#### Bacteria

The following bacterial strains were used as *C. elegans* food: *E. coli* OP50, RNAi-competent *E. coli* rnc14::DTn10 and lacZgA::T7pol camFRT. Bacteria were grown overnight on Luria-Bertani (LB) plates at 37°C from glycerol stocks. The next morning a scoop of the bacterial lawn was inoculated in LB broth with antibiotics (streptomycin for *E. coli* OP50 and grown for 6 h on agitation at 450 g at 37°C (OD600 1.5–2.0). A volume of 300 mL of this bacterial culture was seeded onto 60 mm NGM plates and allowed to dry overnight before larvae or embryos are placed on them.

## Method details

### Dauer quantification and non-dauer/post-dauer separation

#### Fluorescence-based quantification and separation of dauers using the laid-embryo method

For low-throughput experiments, where non dauers were evaluated (Figure 1B, 1D, S1, 6A-D), dauer formation was quantified using the PS8438 strain, which expresses a red fluorescent protein (mCherry) under the control of the dauer-specific *col-183* promoter (*Shih et al.*, 2019). This fluorescence-based marker allows direct identification of dauers under a fluorescence stereoscope. Twenty gravid hermaphrodites were placed on unseeded NGM plates and allowed to lay eggs at room temperature (20– 22°C) for 4 h. Adults were then removed, and the laid embryos were incubated at 27 °C for 45 h (*Karp*, 2018; *Ailion and Thomas*, 2000).

At the end of the incubation period, dauers and non-dauers were counted in triplicate under a fluorescence stereoscope (Nikon SMZ1270) at room temperature. To ensure accurate quantification, each fluorescent dauer was marked on the back of the plate with a colored dot corresponding to its position, and non-dauers were similarly marked using a different-colored pen. A minimum of 30 total animals per plate was required for inclusion in the analysis.

To separate dauer and non-dauer populations, 60–100 red fluorescent dauers were manually picked and transferred to fresh NGM plates seeded with *E. coli* OP50 at room temperature (20–22°C). Non-fluorescent non-dauer (ND) animals were similarly transferred to separate *E. coli* OP50-seeded plates. Both populations were incubated at 20°C. After 24 h, gravid ND hermaphrodites were allowed to lay eggs for 4 h following the standard protocol. Gravid post-dauer (PD) hermaphrodites were obtained 48 h after recovery and subjected to the same procedure. The resulting embryos were used for F1 dauer formation assays.

#### Quantification of pheromone-induced dauer formation

Pheromone crude extraction was performed following Schroeder and Flatt (2014). NGM plates (30 mm) lacking peptone were supplemented with 2% (v/v) pheromone extract and 20 µL of heat-killed *E. coli* OP50. Approximately 15–20 gravid hermaphrodites were placed on each plate and allowed to lay eggs for 4 h (laid-embryo method), after which adults were removed. The resulting embryos were incubated for 48 h at 25°C on pheromone-supplemented plates. Dauer animals were identified by red fluorescence driven by the *col-183* promoter.

#### Temperature-induced dauer formation from bleached embryos

Embryos were isolated from gravid hermaphrodites by standard hypochlorite treatment, washed twice in M9 buffer, and counted in 2 µL aliquots (measured in duplicate) to ensure uniform seeding density. 1,000 embryos were plated per NGM plate and allowed to settle at room temperature for 1 h before transfer to the indicated dauer-inducing condition. This method was used for RNAseq sample preparation (Figure 2), exogenous RNAi and reporter validation (Figure 3A, Figure 4C-D), animals with genetically ablation in specific neurons (Figure 1F), mutant screens (Figure 5A, 5B, 5C) and post-dauer fertility (Figure 6E, 6F).

Plates were incubated at 27°C for 70 h. Dauer animals were selected based on resistance to 1% SDS: plates were washed once into 1.5 mL microcentrifuge tubes with M9 buffer, incubated in 1% SDS for 15 min at room temperature with gentle agitation, and centrifuged to pellet animals. The pellet was washed twice in M9 to remove residual SDS. Dauers were quantified under a stereoscope from at least three 20– 30 µL aliquots per sample. Identification was based on characteristic morphology and confirmed by SDS resistance.

#### Crowding-induced dauer formation (intergenerational assays)

Embryos obtained by hypochlorite treatment from PS8438 gravid adults were plated at high density (12,000 embryos) on *E. coli* OP50-seeded 60 mm NGM plates and incubated at 25 °C for 70 h. Dauers were collected by 1% SDS treatment, washed twice in M9, and counted under a stereoscope from at least three 20–30 µL aliquots per condition (Figure S1).

### Fertility in post-dauer and non-dauer animals

#### Brood size quantification in post-dauer and non-dauer animals and sterility quantification

Brood size was quantified using the PS8438, dauer-specific *col-183* promoter strain, obtained through the laid-embryo and fluorescence-based dauer separation method described above. Following recovery from 27 °C exposure, and for crowding-induced dauers derived from high-density (12,000 embryos) hypochlorite treatment plates, dauers and non-dauers were individually separated by fluorescence.

One day after separation, young adult hermaphrodites of the non-dauer (ND) condition were individually transferred to NGM plates seeded with *E. coli* OP50. Two days after separation, young adult hermaphrodites from post-dauer (PD) condition were similarly transferred to individual *E. coli* OP50 seeded plates.

Hermaphrodites were maintained at 20 °C and allowed to lay eggs. Live progenies were counted every 24 h for four consecutive days. Brood size for each individual was calculated as the total number of live progenies produced during this period. On day 4, plates were inspected to confirm that no additional progeny had emerged. Sterility was confirmed by the absence of embryos or larvae on the plate and by examination of the parental hermaphrodite under a stereoscope to verify the absence of developing embryos in utero. Animals showing no progeny by day 4 were scored as sterile and the percentage of sterile animals was scored in three biological replicates, using Fisher cumulative distribution for statistical analysis.

Each condition was assessed in three biological replicates of 30 animals for temperature (*ND-temp* and *PD-temp*) and 20 animals for crowding (*ND-crowd* and *PD-crowd*). The percentage of sterile animals was calculated based on the total number of animals in each replicate.

#### Brood size/sterility quantification in post-dauer (*PD-temp*) mutant animals

Brood size was estimated for each strain separately, using the hypochlorite treatment as described for the mid-throughput assays. Dauers were obtained by SDS 1% treatment and allowed to recover for 1 day at 20°C. Three biological replicates of 10 individual post-Dauers were transferred to single *E. coli* OP50-seeded plates and maintained at 20 °C. Live progenies were counted every 24 h for four consecutive days, as described above. Brood size was calculated as the total number of live progenies produced during this period. N= 30 individuals. Individuals with 0 progeny were scored as sterile animals and a percentage was calculated.

Mutants analyzed were YY186 *nrde-2(gg91)* II, YY158 *nrde-3(gg66)* X, YY538 *hrde-1* (*tm1200)* III, YY996 *znfx-1*(*gg561*) II, WM27 *rde-1*(*ne219*) V, WM49 *rde-4*(*ne301*) III, NL2098 *rrf-1*(*pk1417*) I, HC271 [ccIs4251 I; *qtIs3 sid-2(qt42)* III; *mIs11* IV], NL3321*sid-1* (*pk3321*) V, RB1815 *vit-3* (*ok2348*)X, MQD2884 *vit-2*(*ok3211*) *vit-1*(*hq532*) X.

### RNA Sequencing

#### Young adults sample preparation

Wild-type *C. elegans* (N2) animals were cultured on NGM plates (60 mm diameter) seeded with *E. coli* OP50. To obtain synchronized populations, embryos were collected via standard hypochlorite treatment and seeded at a density of 1,000 embryos per plate. After 1 hour at RT, animals were incubated at 27 °C for 66 hours (16 plates total). After incubation, plates were brought to room temperature (20°C–22°C), and animals were washed off using M9 buffer. To isolate dauer larvae, the pelleted animals were treated with 1% SDS for 15 minutes with gentle agitation at room temperature. The pellet was then washed twice with M9 buffer to remove residual SDS and plated onto six fresh NGM plates seeded with *E. coli* OP50 to allow recovery. After 24 hours at room temperature, 70–100 synchronized young adults (identified by fully developed vulvae and the absence of embryos in utero) were collected manually and flash-frozen in liquid nitrogen in five biological replicates (quintuplicate). This population was designated *P0-temp*. The remaining *PD-temp* animals were allowed to recover an additional 24 hours until most hermaphrodites became gravid. Embryos were again isolated via hypochlorite treatment and re-exposed to 27°C for 66 hours to obtain the F1 generation. Dauers were isolated via SDS treatment as described above, and 70–100 young adults were collected 24 hours post-recovery and frozen in quintuplicate as the *F1-temp* group.

To assess transgenerational effects, a subset of F1 embryos was allowed to develop at 20°C for one full generation. Four days later, gravid hermaphrodites were collected, bleached, and their embryos were incubated at 27°C for 66 hours. Resulting dauers were selected with 1% SDS, recovered at 20°C, and 70–100 young adults were collected after 24 hours of recovery and stored at -80°C in quintuplicate. This group was designated *F2-temp.* For the pheromone-induced dauer control, N2 embryos (12,000 per plate) were plated at high density on *E. coli* OP50-seeded NGM plates and incubated at 25°C for 66 hours. Animals that entered dauer were subsequently processed using the same SDS isolation, recovery, and young adult collection procedures described above, and stored accordingly for RNA-seq.

#### RNA Extraction

RNA extraction was performed as previously described, with minor modifications (Zhang et al., 2022). Eppendorf frozen worm pellets tubes were transferred to ice at room temperature, and in less than 5 minutes, 1 mL of TRIzol Reagent (Thermo Fisher Scientific) was added to each sample. To facilitate mechanical disruption, 100 µL aliquotes of acid-washed sand (Sigma-Aldrich, catalog no. 274739) was added. Samples were vortexed vigorously for ten minutes.

Phase separation was carried out by adding 200 µL of chloroform, followed by centrifugation at 12,000 × *g* for 15 minutes at 4 °C. The aqueous phase was transferred to a new tube, and RNA was precipitated by the addition of equal volume of isopropanol, then incubated at –20°C for 30 minutes. Pellets were recovered by centrifugation, washed with 75% ethanol, air-dried briefly, and resuspended in nuclease-free water.

Total RNA concentration was quantified using the Qubit RNA HS Assay Kit (Invitrogen, catalog no. Q33224), and RNA integrity was assessed using the Agilent 4150 TapeStation System. Only RNA samples with an RNA Integrity Number (RIN) ≥ 9.0 were used for subsequent Illumina library construction and sequencing.

#### RNA Library Construction and Sequencing

RNA-seq libraries were prepared using standard Illumina protocols in DNase-and RNase-free Eppendorf tubes. Five biological replicates were processed per condition, each in a separate tube. For each sample, 50 ng of total RNA was used to isolate and enrich polyadenylated mRNA using the NEBNext Poly(A) mRNA Magnetic Isolation Module (New England Biolabs, catalog no. E7490L).

Subsequent steps—including RNA fragmentation, first-and second-strand cDNA synthesis, end-repair, and adaptor ligation—were performed using the NEBNext Ultra II RNA Library Prep Kit with Sample Purification Beads (New England Biolabs, catalog no. E7775L), following the manufacturer’s instructions. Adapter ligation was carried out with the NEBNext Multiplex Oligos for Illumina (New England Biolabs, catalog no. E7600A). Libraries were amplified with 14 cycles of PCR.

Library concentrations were quantified using the Qubit dsDNA BR Assay Kit (Invitrogen, catalog no. Q32853). The final libraries were pooled in a single Eppendorf tube, stored at 4°C, and shipped for high-throughput sequencing.

Sequencing was performed on an Illumina NovaSeq 6000 platform, using the XP Workflow with S4 flow cell, 2 × 150 bp paired-end reads, and the 300-cycle v1.5 kit, covering all library pools. The reads were basecalled using Picard Illumina BasecallsToFastq version 2.23.8, with APPLY_EAMSS_FILTER set to false. Following basecalling, the reads were demultiplexed using Pheniqs version 1.1.0 (*Galanti et al.*, 2021) allowing for 1 mismatch in sample index sequences. The entire process was executed using a custom nextflow pipeline, GENEFLOW. (New York University Center for Genomics and System Biology Genomics Core, GitHub Repository. https://github.com/gencorefacility/GENEFLOW; New York University).

### Bioinformatic analysis of *Post Dauer samples*

#### Workflow for mRNA-seq analysis

The following transcriptomic workflow was performed using Galaxy (Galaxy Community, 2024): Initial quality control was performed using FastQC (http://www.bioinformatics.babraham.ac.uk/projects/fastqc) to assess read quality and adapter content. Trimming of low-quality reads and adapter sequences was conducted using Fastp (*Chen et al.*, 2018), with reads discarded if they had a *Phred* quality score <35 or a read length <36 bp, following established criteria (*Gabaldón et al.*, 2020).

Read alignment was performed using HISAT2 (*Kim et al.*, 2015) with default parameters against the *C. elegans* reference genome (WormBase release WS300; updated August 2024). Gene-level quantification was carried out with FeatureCounts (*Liao et al.*, 2014), using the corresponding GTF latest annotation file for *C. elegans* (August 2024).

Differential gene expression analysis was conducted using DESeq2 (*Love et al.*, 2014). Genes with an adjusted P-value (Benjamini–Hochberg correction) <0.001 were considered significantly differentially expressed.

Gene Ontology (GO), Tissue and Phenotype Enrichment Analysis were performed using the enrichment tools available on WormBase (*Angeles-Albores et al.*, 2016; *Angeles-Albores et al.*, 2018), using the set of significantly up-or down-regulated genes per condition filtered by a fold change of 2 and a Q value of 0.001.

#### Clustering analysis (PCA analysis) and heatmap

For Principal component analysis *“rlog”* transformation tool from DESEq2 was included (variance stabilizing transformation) as output. Genes for each sample were ordered by variance, in each comparison, 1000 genes with the highest variance were selected for PCA. Principal Component Scores PC1 and PC2 were selected for visualization Using Prism 10. (Fig 2A, S2A S2B).

Heatmaps were drawn using the seaborn library in python (*Waskom*, 2021). Mean count values were used for each condition by averaging biological replicates. Heatmaps were clustered using correlation distances and average-linkage hierarchical clustering. Expression values (in tpm) log-transformed, centered, and scaled for each gene.

#### RNA interference (RNAi)

Functional validation of candidate genes identified as specifically upregulated in F1 *PD-temp* animals was performed by RNA interference (RNAi) using the bacterial feeding method. The *C. elegans* strain PS8438 was fed *Escherichia coli* HT115(DE3) expressing double-stranded RNA (dsRNA) corresponding to each target gene, using clones obtained from the Ahringer RNAi library (*Fraser et al.*, 2000; *Kamath et al.*, 2003). Animals were maintained on the corresponding RNAi bacteria for two or three consecutive generations, depending on the experimental paradigm described below.

For temperature-induced dauer formation, animals were propagated on RNAi bacteria for two generations at 20°C following the RNAi feeding protocol described above. To initiate the dauer assay, approximately 1,000 gravid adults per condition were subjected to alkaline hypochlorite treatment for 4– 5 min to isolate synchronized embryos. Embryos were pelleted, washed twice with M9 buffer, and approximately 1,000 embryos were seeded onto each of three NGM plates supplemented with carbenicillin and 1 mM IPTG. Plates were left at room temperature for 1 h to allow embryo settlement before incubation at 27°C for 70 h. Dauer larvae were subsequently quantified based on resistance to 1% SDS treatment, as described above.

For starvation-induced dauer formation, animals were maintained on RNAi bacteria for three consecutive generations at 20°C. After depletion of the bacterial food source, plates were left without additional bacteria to induce starvation and maintained at 20°C for two weeks. Dauer larvae were then quantified from three independent plates per condition. The complete experimental procedure, including RNAi propagation and starvation, lasted approximately three weeks. Seven independent biological replicates were performed.

Animals fed *E. coli* HT115(DE3) carrying the empty L4440 vector served as the negative control in all RNAi experiments. The *unc-22* RNAi clone was included as a positive control to verify RNAi efficiency, producing the characteristic twitching phenotype. All RNAi plasmids were purified using a MiniPrep kit and verified by Sanger sequencing with M13-F primers at the Sequencing Unit of Pontificia Universidad Católica de Chile.

#### Statistical analysis

For both the two-generation and three-generation RNAi experiments, statistical comparisons were performed using the Kruskal–Wallis test followed by Dunn’s multiple-comparisons test. Animals fed *E. coli* HT115(DE3) carrying the empty L4440 vector were used as the reference control.

#### Imaging and *vit-3* reporter quantification

Using the strain MQD2775 *vit-3* (hq485[*vit-3::mCherry*]) *vit-2* (crg9070[*vit-2::gfp*]), in which mCherry was knocked into the C-terminus of *vit-3* by CRISPR/Cas9 (*Zhai et al.*, 2022), embryos were subjected to dauer induction at 27°C for 70 h, after which 1% SDS treatment was used to select dauers. Nematodes expressing mCherry were imaged on agar using a Nikon SMZ1270 fluorescence stereomicroscope at 40× magnification, equipped with the appropriate filter cube for RFP fluorescence imaging. All images were acquired using ISO 1600 and a shutter speed of 1/16 s. Images from equivalent developmental stages and experimental conditions were acquired on the same day.

Embryos from animals maintained at 20°C, as well as F1 embryos derived from PD animals, were scored. L2 animals from the P0 and F1 generations were selected 24 h post-hatching at 27°C. High-temperature-induced dauers were scored immediately after SDS treatment, 70 h after incubation at 27°C. At least 40 embryos, 28 L2 animals, and 20 dauers were quantified for each condition.

Using ImageJ, the complete body of each individual was selected, and the average pixel intensity was measured and recorded as arbitrary fluorescence units. Background fluorescence was also measured for each image. Background fluorescence values were subtracted from the corresponding individual fluorescence measurements, and the resulting corrected values were used for quantification and analysis.

#### Survival assay after HID

Using the blacked embryo protocol, assessment of survival after dauer induction of wild type and mutant embryos growing for 70 hours at 27°C. The number of alive animals was scored in three technical replicates each in an aliquot of 20uL each, the mean number of animals was then multiplied for the dilution factor. The mean total number of animals alive compared to the initial number of eggs initially counted which was around 1000. N = 5 biological replicates

### Quantification and statistical analysis

#### Experimental replicas and statistical evaluation

All experiments were done at least in three biological replicas or more (stated in methods and in Stats dataset), started on different days and from different starting plates. Each biological replica contained a triplicate (three technical replicas). Data in graphs are represented as mean ± SEM. Figure with asterisks represent p-values as follows: ∗<0.0332, ∗∗<0.0021, ∗∗∗<0.0002, ∗∗∗∗<0.0001. Statistical evaluation was done by a one or two-way ANOVA with post-hoc analyses, in the case of sterility rates, statistical significance was assessed using Fisher’s exact test on cumulative counts. Results of all experiments in the manuscript and statistical tests including descriptive statistics are detailed in **Dataset 4 and 5** respectively.

#### Criteria for data exclusion

We excluded experimental replicas when there was contamination with unwanted bacteria or fungi on the nematode plates, or when bacteria had been almost or completely consumed.

#### Temperature variation of incubators

Incubation temperatures were monitored throughout all experiments. For assays performed at 25°C, the temperature inside the incubator was measured at the end of each experiment, yielding an average of 25.5 ± 0.3°C. Similarly, for assays conducted at 27 °C, temperature monitoring revealed slightly higher values, with an average of 27.5 ± 0.4°C across experiments.

## Acknowledgments

We thank Erik C. Andersen for his support of the transcriptomic experiments, including providing laboratory resources and funding, and for his valuable input on data analysis and interpretation. We are also grateful to Cori Bargmann for critical reading of the manuscript. Some strains were provided by the CGC, which is funded by the NIH Office of Research Infrastructure Programs (P40OD010440). This work was funded by Fondecyt Grant 1220650 and the Millennium Scientific Initiative ICM-ANID (ICN26-30, CINV) to AC. ER was supported by doctoral fellowship 21211299 from the National Agency for Research and Development (ANID) and by a Beca de Extensión DNUV from the University of Valparaíso. JL is supported by a grant from the Chan Zuckerberg Initiative to C. Bargmann.

## Author contributions

Conceptualization ER and AC, Methodology ER, JL, and AC, Investigation ER, JL, and AC, Writing-Original Draft ER and AC, Writing-Review and Editing ER, JL and AC, Funding Acquisition AC.

## Notes

### Competing Interest Statement

The authors have declared no competing interest.

